# Investigating causality between liability to ADHD and substance use, and liability to substance use and ADHD risk, using Mendelian randomization

**DOI:** 10.1101/524769

**Authors:** Jorien L Treur, Ditte Demontis, George Davey Smith, Hannah Sallis, Tom G Richardson, Reinout W Wiers, Anders D Børglum, Karin JH Verweij, Marcus R Munafò

## Abstract

Attention-deficit hyperactivity disorder (ADHD) has consistently been associated with substance use, but the nature of this association is not fully understood. To inform intervention development and public health messages, a vital question is whether there are causal pathways from ADHD to substance use and/or vice versa. We applied bidirectional Mendelian randomization, using summary-level data from the largest available genome-wide association studies (GWAS) on ADHD, smoking (initiation, cigarettes/day, cessation, and a compound measure of lifetime smoking), alcohol use (drinks/week, alcohol problems, and alcohol dependence), cannabis use (initiation) and coffee consumption (cups/day). Genetic variants robustly associated with the ‘exposure’ were selected as instruments and identified in the ‘outcome’ GWAS. Effect estimates from individual genetic variants were combined with inverse-variance weighted regression and five sensitivity analyses (weighted median, weighted mode, MR-Egger, generalized summary-data-based-MR, and Steiger filtering). We found evidence that liability to ADHD increases likelihood of smoking initiation and heaviness of smoking among smokers, decreases likelihood of smoking cessation, and increases likelihood of cannabis initiation. There was weak evidence that liability to ADHD increases alcohol dependence risk, but not drinks/week or alcohol problems. In the other direction, there was weak evidence that smoking initiation increases ADHD risk, but follow-up analyses suggested a high probability of horizontal pleiotropy. There was no clear evidence of causal pathways between ADHD and coffee consumption. Our findings corroborate epidemiological evidence, suggesting causal pathways from liability to ADHD to smoking, cannabis use, and, tentatively, alcohol dependence. Further work is needed to explore the exact mechanisms mediating these causal effects.

## Introduction

Individuals who have been diagnosed with attention deficit hyperactivity disorder (ADHD) are more likely to be (heavy) substance users compared to those without ADHD^1^. Around 5.9-7.1% of children and adolescents and 5.0% of adults are thought to meet the diagnostic criteria for ADHD^2^, and genetic studies support the notion that a clinical diagnosis represents the extreme end of a continuum of impulsivity and/or attention problems in the general population^3, 4^. Both ADHD diagnosis and higher levels of impulsivity and attention problems are associated with higher levels of cigarette smoking^5, 6^, cannabis use^7, 8^, alcohol use^7, 9^ and caffeine consumption^10, 11^. The exact nature of these associations is not fully understood, which hampers the development of evidence-based interventions and public health messages.

Several explanations have been posited as to why ADHD and substance use are correlated. First, there are risk factors that increase susceptibility to both. These could be environmental factors that have been shown to be risk factors for ADHD and substance use, such as trauma exposure or other adverse early life events^12, 13^, or these could be genetic influences with pleiotropic effects on both ADHD and substance use. Family studies have shown that ADHD and substance use are moderately to highly heritable, and indicate shared genetic risk factors^14, 15^. Overlap in genetic risk has also been examined in recent genome-wide association studies (GWAS) of ADHD and substance use^4, 16–19^. Substantial genetic correlations were found for ADHD with ever versus never smoking (r_g_=0.48, *p*=4.3e-16), number of cigarettes smoked per day (r_g_=0.45, *p*=1.1e-05), alcohol dependence (r_g_=0.44, *p*=4.2e-06), and cannabis initiation (r_g_=0.16, p=1.5e-04), pointing to a common neurobiological aetiology. This is consistent with research indicating that cognitive deficits such as impaired response inhibition and working memory are important features of both ADHD and substance abuse^20, 21^, and that both ADHD and substance abuse can be considered forms of externalizing disorders^22^.

While a partial common neurobiological aetiology to ADHD and substance use is therefore likely, environmental and genetic correlation could also (partly) reflect causal effects of one on the other. If variable X causes variable Y, it follows that any environmental or genetic risk factor causing variable X will also be associated (indirectly) with variable Y. The current literature has mostly focused on causal pathways from ADHD to substance use, with longitudinal cohort studies showing that externalizing symptoms in early adolescence predict onset of smoking and faster progression to daily smoking, and that ADHD medication reduces early onset smoking and alleviates smoking withdrawal^5^. For alcohol and cannabis the evidence is less clear, with some studies finding that ADHD symptoms only predict their use in girls^23^, and a recent twin study reporting no relation between ADHD symptoms and alcohol or cannabis use^14^. For caffeine, a relatively small longitudinal study (*n*=144) suggested reciprocal effects between caffeine consumption and ADHD symptoms during adolescence^10^.

There is tentative evidence that there may be causal effects in the other direction (i.e., substance use leading to an increase in ADHD symptoms)^24, 25^. In monozygotic twin pairs discordant for smoking, the smoking twin scored higher on attention problems – a difference which only appeared after smoking was initiated^24^. For cannabis use the evidence is mixed. Low to moderate cannabis use in adolescents seems to lead to a small increase in attention and academic problems, which disappears following sustained abstinence^25^. However, there is no indication that cannabis use exacerbates ADHD-related brain alterations^26^. With regards to alcohol use, binge-pattern exposure during development has been shown to cause attention deficits in mice^27^, but there is no clear evidence for such effects in humans.

It is difficult to fully unravel the nature of the association between ADHD and substance use with observational data because of bias due to (unmeasured) confounding and reverse causality (i.e., the outcome affecting the exposure). Mendelian randomization (MR) is a method to infer causality which has recently gained much popularity. MR uses genetic variants robustly associated with an exposure variable as an instrument to test causal effects on an outcome variable^28, 29^. Because genes are transmitted from parents to offspring randomly, genetic variants that are inherited for a trait (e.g., ADHD) should not be associated with confounders such as social-economic status. By using genetic variants as instrumental variables it is therefore possible to obtain less biased results.

So far, one MR study found evidence for a causal effect of alcohol use on attention problems and aggression in adolescents (but not on delinquency, anxiety, or depression)^30^. Two other studies provided evidence that genetic liability to ADHD, as well as higher extraversion, has a causal effect on smoking initiation^31, 32^. A recent MR study found that liability to ADHD leads to a higher risk of cannabis initiation, but these analyses were based on summary-level data of a cannabis GWAS which has recently been updated (with a much larger sample size – n=184,765 instead of n=32,330). Moreover, potential causal effects in the reverse direction were not adequately tested given that the authors included all ADHD cases instead of just those diagnosed in adulthood^33^. Overall, existing MR studies are limited in that they have primarily tested unidirectional effects only, included a narrow focus on one specific substance use behaviour, and/or had limited statistical power.

We therefore performed bidirectional MR using summary-level data of the largest available GWAS, investigating causal effects between liability to ADHD and a broad spectrum of substance use phenotypes. We applied five different sensitivity analyses more robust to potential violation of the MR assumptions. Throughout the manuscript, we refer to *‘liability to’* a particular exposure (e.g. liability to ADHD). This is because the exposure estimates and the outcome estimates for our analyses come from separate samples, and it is not possible to determine whether or not the individuals in the outcome sample have actually experienced a particular exposure (e.g. an ADHD diagnosis).

## Materials and methods

### Mendelian randomization

The rationale behind MR is that the random assortment of genetic variants creates subgroups in the population which roughly mimic treatment groups from a randomized controlled trial. Outcomes are compared between individuals in the ‘high genetic risk group’ and those in the ‘low genetic risk group’ for a proposed exposure variable. The method rests on three important assumptions, namely that the genetic variants used as instruments: 1) strongly predict the exposure variable – typically, the selected variants have been genome-wide significantly associated (*p*<5e-08) with the exposure and replicated, 2) are independent of confounding variables, and 3) do not affect the outcome through an independent pathway, other than possible causal effects via the exposure (**Figure 1a**). A potential threat to MR is horizontal pleiotropy, where the genetic variant used as an instrument *directly* affects vulnerability to multiple phenotypes. This could lead to violation of MR assumptions 2 and 3. To assess whether MR assumptions may have been violated, we conducted various sensitivity analyses described below.

**Figure 1.**
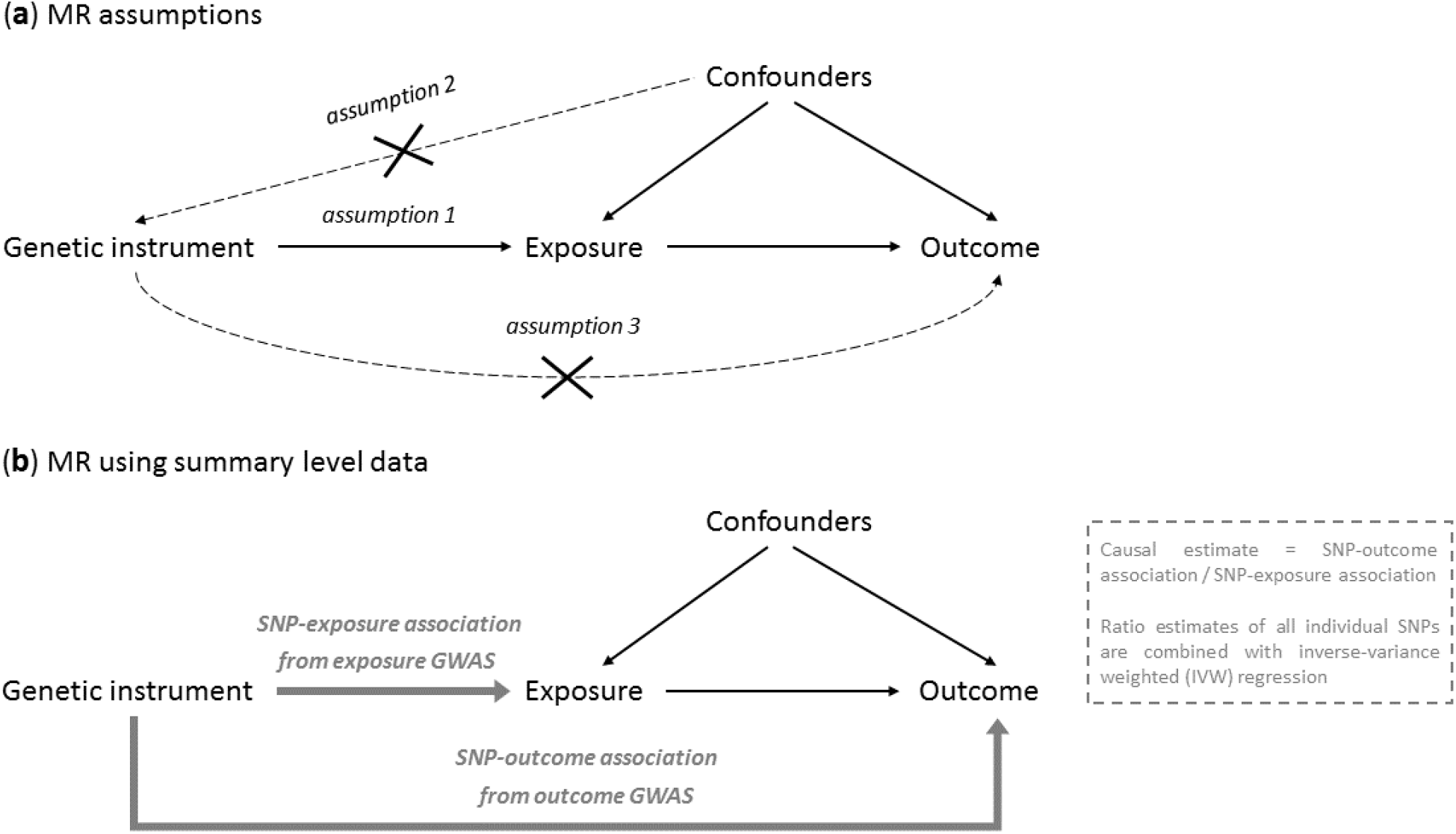
**a)** Illustration of the Mendelian randomization (MR) framework and its main assumptions that the instrument is associated with the exposure (1), the instrument is not associated with (un)measured confounders (2), and the instrument does not influence the outcome other than through the exposure (3). **b)** Illustration of the MR design when using summary level data and the SNP-exposure association and SNP-outcome association are taken from two separate GWASs (also known as ‘two-sample MR’).

We applied MR using summary level data (sometimes known as ‘two-sample MR’), which uses effect estimates of genetic variants (SNPs) from large GWAS that have been performed previously. In this approach, the SNP-exposure association and the SNP-outcome association estimates are taken from two separate GWAS (**Figure 1b**). A major strength of this design is that it takes advantage of large, well-powered GWAS, without the need to have information on both the exposure and the outcome in one single sample. An additional assumption of this method is that the SNPs identified as instruments based on their effect estimates in the exposure GWAS also predict that exposure variable in the outcome GWAS – this cannot be directly tested. To estimate the causal effect of the exposure on the outcome, the SNP-outcome association is divided by the SNP-exposure association for each SNP. The main MR result is obtained by combining these ratios into an overall estimate of causal effect using inverse-variance weighted (IVW) fixed-effect meta-analysis (**Figure 1b**)^29^.

### Mendelian randomization versus other causally informative designs

MR is inherently different from other causally informative designs such as twin or family studies. While these methods use *a priori* knowledge of genetic relatedness between family members to explain variation in a particular phenotype, MR exploits directly measured genotype in (usually) unrelated individuals. Whereas twin and family studies aim to correct for genetic (and shared environmental) differences in order to infer causality, MR exploits the genetic component by using it as an instrument for causal inference. A comprehensive review comparing all methods that use genetic data to strengthen causal inference is available elsewhere^34^.

### Data

Summary-level data of large GWAS were obtained for ADHD (clinically diagnosed versus controls, *n*=55,374^4^), smoking (initiation (ever regularly smoked or ≥100 cigarettes during lifetime), *n*=1,232,091; cigarettes per day, *n*=337,334; cessation (former versus current smokers), *n*=547,219^17^; lifetime smoking, *n*=463,003^35^), alcohol use (drinks per week, *n*=941,280^17^; alcohol problems (AUDIT total score: Alcohol Use Disorders Identification Test), *n*=121,604^19^; alcohol dependence (clinically diagnosed versus controls), *n*=46,568^36^), cannabis use (initiation (ever used during lifetime), *n*=162,082^16^) and coffee consumption (cups per day, *n*=91,462^18^). When smoking initiation, cigarettes per day, smoking cessation or alcohol drinks per week was the outcome in the MR analysis, data of one of the included cohorts, 23andMe, were not available, resulting in sample sizes of *n*=632,783, *n*=263,954, *n*=312,821 and *n*= 537,341, respectively.

Lifetime smoking is a compound variable that captures smoking initiation, duration, heaviness and cessation, across mid- to late-adulthood. As ADHD onset is expected to occur (long) before mid- to late-adulthood, lifetime smoking was not appropriate to use as an exposure and was only used as an outcome. In addition, cigarettes per day and smoking cessation could not be used as exposures because the GWAS these are based on were performed in (former) smokers only. To perform an MR analysis with genetic variants for cigarettes per day and smoking cessation as instruments, the outcome GWAS (in this case ADHD) would have to be stratified on smoking status, which was not possible with the summary data we used.

When testing causal effects of liability to ADHD on substance use, summary statistics from the complete ADHD GWAS containing child, adolescent, and adult data were used. When testing causal effects of substance use on ADHD, only adult data (ADHD diagnosed >18 years) were used (*n*=15,548) to ensure a plausible temporal sequence of a potential causal effect (i.e., substance use cannot logically have a causal effect on ADHD diagnosed in childhood). This is crucial given that ADHD is generally a child-onset disorder and the onset of substance use is typically during adolescence or early adulthood. What we aim to test here is whether substance use causes later development of or exacerbating of ADHD symptoms (resulting in a diagnosis at adult age). There was no sample overlap of the ADHD GWAS with the smoking, alcohol and coffee GWAS. Between the ADHD and the cannabis initiation GWAS there was very minimal overlap (<3%).

### Main analysis

To assess causal effects of liability to ADHD on substance use, we identified independent single nucleotide polymorphisms (SNPs) that reached genome-wide significance (*p*<5e-08) in the ADHD GWAS to use as genetic instruments. These same SNPs were then identified in the substance use GWAS. To assess causal effects in the other direction, we identified independent genome-wide significant SNPs in the different substance use GWAS as genetic instruments, and then identified those different sets of SNPs in the ADHD GWAS. The analyses were conducted in R, using the two-sample MR package of MR-Base, a database and analytical platform^37^.

### Sensitivity analyses

Besides IVW (**Figure 1b**), five additional MR methods were applied. Each of these five methods can provide an unbiased estimate of the true causal effect, provided that certain assumptions are met. While it is not possible to know which of these methods’ assumptions actually hold, examining the combined results of all methods allows us to assess the robustness of a causal finding. This practice of using multiple MR methods to triangulate evidence has become increasingly important now that MR has moved beyond more biological phenotypes (e.g., LDL-cholesterol) and is increasingly used in the context of complex traits such as ADHD and substance use, were detailed knowledge of the exact biological function of the associated genes is lacking^38^.

First, we used weighted median regression, which provides an unbiased estimate of the causal effect, even if <50% of the weight of the genetic instrument comes from invalid instruments^39^. Second, we used weighted mode regression, which provides unbiased results as long as the causal effect estimate that is most common among the included SNPs comes from valid instruments and is thus consistent with the true causal effect^40^. Third, we used MR-Egger regression, which provides an unbiased estimate of the causal effect provided that the strength of the genetic instrument (association between the SNP and the exposure) does not correlate with the effect that same instrument has on the outcome. This ‘InSIDE assumption’ (Instrument Strength Independent of Direct Effect) is a weaker assumption than the assumption of no pleiotropy^41^. However, MR-Egger does rely on the NOME (NO Measurement Error) assumption, and if this is violated its results may be biased. Violation of the NOME assumption can be assessed by the I^2^ statistic. An I^2^ value below 0.9 indicates considerable risk of bias, which may still be corrected for with MR-Egger simulation extrapolation (SIMEX). An I^2^ value below 0.6 means that MR-Egger results (even with SIMEX) are unreliable. We report MR-Egger results when I^2^>0.9, report MR-Egger SIMEX results when I^2^=0.6-0.9, and do not report MR-Egger results when I^2^<0.6^42^. Fourth, we used generalised summary-data-based Mendelian randomization (GSMR)^43^. This method achieves higher statistical power than other MR methods by taking into account very low levels of linkage disequilibrium (LD) between the included SNPs. GSMR includes a filtering step which identifies and removes SNPs considered outliers based on their effect size (HEIDI-filtering). MR-Egger and GSMR were applied only when the genetic instruments contained 10 or more SNPs. Fifth, we used Steiger filtering, which computes the amount of variance each SNP explains in the exposure and in the outcome variable. In case of a true causal effect of the exposure on the outcome, a SNP used as an instrument should be more predictive of the exposure than the outcome. If not (i.e., the SNP is more predictive of the outcome than the exposure) it might imply reverse causation^44^. Steiger filtering was used to exclude all SNPs that were more predictive of the outcome than the exposure, after which MR analyses were repeated. LCV (Latent Causal Variable model) is a recent method with the potential to distinguish genetic correlation from causation^45^. While we conducted LCV analyses, we report these in the supplemental material only because: 1) we aim to explicitly test bidirectional causality, which LCV does not allow, and 2) for cigarettes smoked per day, smoking cessation, and lifetime smoking LCV is not appropriate because we intended to only use them as outcome variables, and with LCV it is not possible to indicate which trait is the exposure or outcome.

For an additional indication of the robustness of our findings we inspected the Cochran’s Q statistic, which provides an estimate of heterogeneity between the effects of the individual genetic variants^46^, and performed leave-one-out analyses, repeating the IVW analysis after removing each of the SNPs one at a time^37^.

### Defining strength of evidence

We did not explicitly correct for multiple testing to avoid judging the evidence based simply on an arbitrary threshold. Instead, we interpret the evidence by looking at both the effect size and statistical evidence for the main IVW result, combined with how consistent the results of the sensitivity analyses are across multiple MR methods. Due to their stricter assumptions, the sensitivity analyses have lower statistical power to identify a true causal effect. Thus, when the effect sizes of the sensitivity analyses are of similar magnitude and direction, this supports a causal interpretation, even if the statistical evidence for an individual analytical approach is weaker than in the IVW analysis.

## Results

We found evidence for causal effects of liability to ADHD on smoking initiation (IVW beta=0.07, 95% CI=0.03 to 0.11, *p*=1.7e-05), cigarettes smoked per day (IVW beta=0.04, 95% CI=0.02 to 0.06, *p*=0.006), smoking cessation (IVW beta=−0.03, 95% CI=−0.05 to −0.01, *p*=0.005) and lifetime smoking (IVW beta=0.07, 95% CI=0.06 to 0.14, *p*=1.4e-07). The weighted median and weighted mode sensitivity analyses confirmed these findings, albeit with slightly weaker statistical evidence for the latter (**Table 1**). For smoking initiation, MR-Egger did not show clear evidence for a causal effect, but this may have been due to a lack of statistical power^41^. The Egger intercept did not indicate horizontal pleiotropy (intercept=0.01, 95% CI=−0.01 to 0.02, *p*=0.41, **Supplementary Table 2**). For cigarettes smoked per day and smoking cessation MR-Egger also did not confirm the IVW findings, with weak evidence for horizontal pleiotropy (Egger intercept=0.01, 95% CI=0.00 to 0.02, *p*=0.068 and intercept=0.01, 95% CI=0.00 to 0.01, *p*=0.089, respectively). GSMR could not be performed because there were too few SNPs (<10). Steiger filtering showed that – with the exception of one SNP in the ADHD risk to smoking initiation analysis – all SNPs were more predictive of the exposure than of the outcome. Cochran’s Q statistic indicated heterogeneity of the effects of the included variants for the ADHD liability to smoking initiation and ADHD liability to lifetime smoking analyses (**Supplementary Table 3**; Q=34.44, *p*=7.5e-05 and Q=47.73, *p*=2.9e-07, respectively), while leave-one-out analyses gave no indication that the overall causal effect was driven by a particular SNP (**Supplementary Figure 1**).

**Table 1.** Results of the Mendelian randomization analyses using summary level data from liability to ADHD to substance use risk including IVW estimates and four sensitivity analyses: weighted median, weighted mode, MR-Egger, and GSMR (generalized summary-data-based Mendelian randomization).

| Exposure | Outcome | IVW |  |  |  |  | Weighted median |  |  |  |  | Weighted mode |  |  |  |  | MR-Egger |  |  |  |  | GSMR |  |
| --- | --- | --- | --- | --- | --- | --- | --- | --- | --- | --- | --- | --- | --- | --- | --- | --- | --- | --- | --- | --- | --- | --- | --- |
|  |  | *n*<br>SNPs | beta | OR | 95% CI | *p* | beta | OR | 95% CI | *p* | beta | OR | 95% CI | *p* | beta | OR | 95% CI | *p* | *n*<br>SNPs* | beta | OR | 95% CI | *p* |
| ADHD | Smoking initiation | 10 | 0.07 |  | 0.03 to 0.11 | 1.7e-05 | 0.05 |  | 0.03 to 0.07 | 4.2e-05 | 0.05 |  | 0.03 to 0.07 | 0.010 | 0.01 |  | -0.13 to 0.15 | 0.937 | 8 | *n.a.* |  | *n.a.* | *n.a.* |
| ADHD | Cigarettes / day | 10 | 0.04 |  | 0.02 to 0.06 | 0.006 | 0.05 |  | 0.01 to 0.09 | 0.004 | 0.05 |  | -0.01 to 0.11 | 0.089 | -0.11 |  | -0.25 to 0.03 | 0.127 | 8 | *n.a.* |  | *n.a.* | *n.a.* |
| ADHD | Smoking cessation | 11 | -0.03 |  | -0.05 to -0.01 | 0.005 | -0.03 |  | -0.05 to -0.01 | 0.026 | -0.03 |  | -0.07 to 0.01 | 0.215 | 0.06 |  | -0.04 to 0.16 | 0.255 | 9 | *n.a.* |  | *n.a.* | *n.a.* |
| ADHD | Lifetime smoking | 10 | 0.10 |  | 0.06 to 0.14 | 8.8e-08 | 0.09 |  | 0.05 to 0.13 | 1.6e-09 | 0.10 |  | 0.06 to 0.14 | 0.003 | *n.a.* |  | *n.a.* | *n.a.* | 9 | *n.a.* |  | *n.a.* | *n.a.* |
| ADHD | Alcohol drinks / week | 10 | -0.01 |  | -0.05 to 0.03 | 0.741 | 0.02 |  | 0.00 to 0.04 | 0.150 | 0.02 |  | 0.00 to 0.04 | 0.153 | 0.08 |  | 0.06 to 0.10 | 0.468 | 7 | *n.a.* |  | *n.a.* | *n.a.* |
| ADHD | Alcohol problems | 10 | 0.01 |  | -0.01 to 0.03 | 0.234 | 0.00 |  | -0.02 to 0.02 | 0.729 | 0.00 |  | -0.02 to 0.02 | 0.815 | *n.a.* |  | *n.a.* | *n.a.* | 8 | *n.a.* |  | *n.a.* | *n.a.* |
| ADHD | Alcohol dependence | 12 | 0.07 | 1.07 | 1.01 to 1.14 | 0.030 | 0.06 | 1.06 | 0.96 to 1.17 | 0.230 | 0.07 | 1.07 | 0.93 to 1.23 | 0.378 | -0.16 | 0.85 | 0.60 to 1.21 | 0.407 | 10 | 0.06 | 1.06 | 0.98 to 1.15 | 0.094 |
| ADHD | Cannabis initiation | 10 | 0.12 | 1.13 | 1.02 to 1.25 | 0.010 | 0.17 | 1.19 | 1.05 to 1.34 | 0.001 | 0.19 | 1.21 | 0.97 to 1.52 | 0.107 | *n.a.* | *n.a.* | *n.a.* | *n.a.* | 10 | 0.11 | 1.12 | 1.03 to 1.21 | 0.004 |
| ADHD | Cups of coffee / day | 9 | 0.03 |  | -0.03 to 0.09 | 0.322 | 0.04 |  | -0.04 to 0.12 | 0.331 | 0.05 |  | -0.09 to 0.19 | 0.458 | *n.a.* |  | *n.a.* | *n.a.* | 9 | *n.a.* |  | *n.a.* | *n.a.* |

*n* SNPs = number of SNPs included in the genetic instrument. SE = standard error of the beta. Note that the dichotomous variables smoking initiation and smoking cessation were rescaled in the original GWAS such that its unit is a standard deviation increase in prevalence<sup>17</sup>. The beta coefficients in this table represent the change in outcome per 2.72-fold increase in the prevalence of ADHD diagnosis (due to the log odds nature of the ADHD GWAS data). For MR-Egger; when $I^2$ was 0.6-0.9, a SIMEX correction was applied, while estimates were not reported at all when $I^2$ was <0.6. *n.a.*: the number of SNPs available for the analysis was too low, or, in the case of MR-Egger, $I^2$ was <0.6.
\*Number of SNPs left after the HEIDI filtering step which is part of GSMR.

There was also considerable evidence that liability to ADHD causally increases risk of cannabis use initiation (IVW OR=1.13, 95% CI=1.02 to 1.25, *p*=0.010). Weighted median, weighted mode and GSMR confirmed this finding, but with (slightly) weaker statistical evidence. MR-Egger was not reported due to a low I^2^ value (**Supplementary Table 4**). Steiger filtering did not identify any SNPs more predictive of the outcome than of the exposure. There was weak evidence for heterogeneity in SNP effects for the ADHD liability to cannabis initiation analysis (Q=15.90, *p*=0.069). Leave-one-out analyses did not suggest any individual SNPs were driving the overall effect.

There was no clear evidence for a causal effect of liability to ADHD on alcohol drinks per week, alcohol problems or coffee consumption. While there was some weak evidence that liability to ADHD causally influences alcohol dependence (IVW OR=1.07, 95% CI=1.01 to 1.14, *p*=0.030), this effect was not consistent across the sensitivity analyses. However, when we repeated these analyses using alcohol intake *frequency* as the outcome measure in UK Biobank only – one of the cohorts included in the much larger GWAS sample the main analyses were based on – there was evidence for a causal effect reflecting increased risk (IVW beta=0.22, 95% CI=0.04 to 0.40, *p*=0.013, **Supplemental Table 5**). This is in line with recent findings that different alcohol use behaviours can show distinct (directions of) genetic associations^47^.

In the other direction, we found strong evidence for causal effects of liability to smoking initiation on ADHD risk (IVW OR=3.72, 95% CI=3.10 to 4.44, *p*=2.9e-51). Weighted median, weighted mode, MR-Egger, and GSMR sensitivity analyses indicated similarly strong evidence, albeit with smaller effect sizes (**Table 2**). The Egger intercept did not indicate horizontal pleiotropy (intercept=0.01, 95% CI=−0.01 to 0.03, *p*=0.37). However, for this relationship the I^2^ value was low – 0.60 (**Supplementary Table 4**) – indicating that MR-Egger was not reliable. Furthermore, Steiger filtering revealed that only 265 of the 346 smoking initiation SNPs (77%) were more predictive of the exposure, smoking, than of the outcome, ADHD. When repeating the IVW and sensitivity analyses with these SNPs only, the evidence for a causal effect was still strong, but effect sizes were attenuated (**Supplementary Table 6**). Cochran’s Q statistic provided no clear evidence for heterogeneity for the liability to smoking initiation to ADHD risk analysis (Q=373.84, *p*=0.14) and leave-one-out analyses did not indicate that the overall effects were driven by a single SNP. As an additional sensitivity test, we repeated the smoking initiation – ADHD analyses using ADHD symptoms in childhood only, with one of the replication samples of the original GWAS paper (<13 years; n=17,666^48^). The degree to which smoking initiation SNPs predict ADHD childhood symptoms in such an MR analysis could reflect horizontal pleiotropy, since most individuals in this age group will not have begun to smoke yet. We found strong evidence for a causal effect (IVW beta=0.28, 95% CI=0.17 to 0.39; **Supplementary Table 7**) – although these effect estimates and the statistical evidence were weaker than in the original analyses that restricted to adults. Together with the results of Steiger filtering, this indicates that the increasing effect of smoking initiation on ADHD risk is, at least in part, due to horizontal pleiotropy.

**Table 2.**
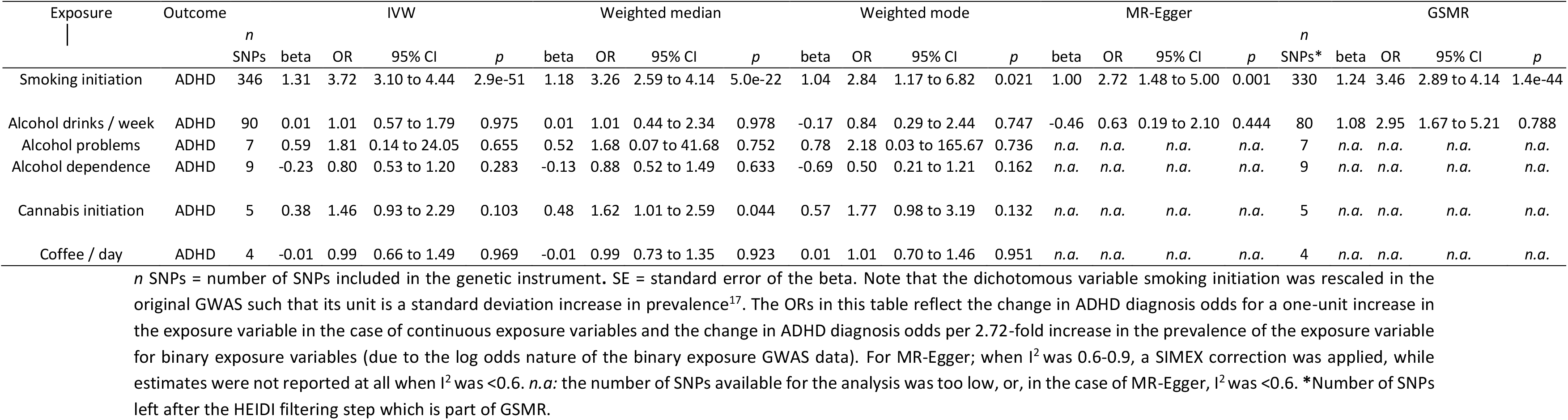
Results of the Mendelian randomization analyses using summary level data from liability to substance use to *adult* ADHD risk (diagnosis received after age 18) including IVW estimates and four sensitivity analyses: weighted median, weighted mode, MR-Egger, and GSMR (generalized summary-level-data based Mendelian randomization).

There was no clear evidence for a causal effect of liability to cannabis use initiation, alcohol use, or coffee consumption on ADHD risk.

The results of LCV analyses indicated that smoking initiation and alcohol dependence are genetically causal for ADHD, while for all other relationships there was no clear evidence of causal effects (**Supplementary Table 8**).

## Discussion

We find evidence, using Mendelian randomization analyses of summary-level data, for causal effects of liability to ADHD on substance use risk, such that it increases the odds of initiating smoking, smoking more cigarettes per day among smokers, and finding it more difficult to quit, as well increasing the odds of initiating cannabis use. There was some indication that liability to ADHD increases alcohol dependence risk, but evidence for that was weak. In the other direction there was weak evidence that liability to smoking initiation increases (adult) ADHD risk. There was no clear evidence of causal effects between liability to ADHD and coffee consumption.

Our findings complement and confirm a large body of observational literature suggesting that individuals diagnosed with ADHD are at a higher risk of initiating smoking, transitioning into regular smoking, and being less able to quit^5^. We also provide evidence for a causal effect of liability to ADHD on risk of cannabis use, for which the literature has so far been inconclusive^14, 23^. While previous observational studies may have been biased by (unmeasured) confounding, our approach of using genetic variants as instrumental variables is more robust to confounding and reverse causality. We were not able to identify the exact mechanism of causation, but it seems plausible that higher levels of impulsivity may lead individuals with ADHD liability to try out cigarettes or cannabis without considering their possible negative consequences^5, 49^. Another potential mechanism is ‘self-medication’, whereby a substance is used because of its (real or perceived) positive effects on ADHD symptomatology – even though such effects might not actually exist^50^.

Interestingly, there was also evidence for causal effects of liability to smoking on ADHD risk. This is in line with previous literature indicating that smoking can have detrimental, long-term effects on attention^24^. It has been hypothesized that nicotine inhaled through cigarette smoke can affect the developing prefrontal cortex – involved in attention and impulse control – during adolescence^51^. It is important to note, however, that the evidence we found for causal effects of smoking on ADHD risk was much less robust than it was in the other direction. First of all, we were not able to test causal effects of smoking heaviness or smoking cessation on ADHD, which would have provided more compelling evidence. Second, a considerable portion (23%) of the SNPs used as an instrument for smoking initiation were in fact more predictive of the outcome, ADHD, implying reverse causation. There is extensive research showing that genetic influences on smoking initiation are mediated via impulsivity-related traits^5^. This was confirmed by our findings that the genetic instrument for smoking initiation also showed strong evidence for a causal effect on ADHD symptoms in children <13 years (who would not yet have started smoking). Another important point is that for the analyses of substance use to ADHD, we used adult diagnosed ADHD as the outcome. This strengthened our approach by ensuring the appropriate temporal sequence for a causal effect in this direction. However, it might be that individuals with adult diagnosed ADHD differ from those who were diagnosed during childhood. A recent study assessed the neurodevelopmental profile of individuals diagnosed with ADHD in adulthood, and found that they did not have a typical profile of neurodevelopmental impairment^52^. Our results should therefore be replicated using other, continuous measures of ADHD symptoms in adulthood. Preferably these would be more ‘proximal’ measures of attention problems and impulsivity, obtained through cognitive performance tasks or (functional) brain imaging.

We found some weak evidence for a causal effect of liability to ADHD on alcohol dependence risk (based on a DSM-diagnosis), but no clear evidence for causal effects of liability to ADHD on drinks per week or alcohol problems (based on the AUDIT self-report survey). These findings are of particular interest because current evidence on the mechanisms underlying associations between ADHD and alcohol use is inconclusive^14, 23^. Given the very large and powerful genetic datasets that our analyses are based on, one would expect that a strong causal effect of ADHD on alcohol use would be convincingly shown, which was not the case given the weak evidence. The fact that there was some indication of causality from ADHD liability to alcohol dependence risk, but not for the other two alcohol measures, weakens the evidence further. However, it might be that ADHD liability only affects serious manifestations of alcohol abuse – such that it is clinically diagnosed – but not self-reported consumption. Of all the included GWAS data sets included in our study, alcohol dependence was based on the smallest sample size (n=46,568), and so it would be good to attempt replication of this finding when bigger samples become available. There was no clear evidence for causal effects between liability to ADHD and coffee consumption, which would indicate that observational correlations are the result of shared risk factors rather than causality.

Important strengths of this study include the very large and recent samples that the analyses are based on, the variety of different substance use phenotypes that were included, and the use of multiple sensitivity analyses that each rely on distinctly different assumptions. However, there are also limitations to consider. First, the genetic instruments used in MR may vary in their strength (i.e., the amount of variance in the exposure variable that they explain). Stronger instruments are more likely to identify a causal effect, which in theory could explain why there was reasonable evidence for causality for some relationships (e.g., smoking to ADHD), but not for others (e.g., alcohol to ADHD). When looking at the predictive power of the instruments, the differences were modest – for ADHD all SNPs included in the instrumental variable combined explained 0.5-0.7% of the variance, for smoking initiation 2.4%, for cannabis initiation 0.2%, for alcohol drinks per week 1.1%, for alcohol problems 0.3%, for alcohol dependence 0.2%, and for coffee consumption 0.6% (the formula to compute these numbers is described elsewhere^16^). However, the power of these instruments to pick up effects on the outcomes also depends on the sample size of the outcome samples. Second, we were not able to apply all sensitivity analyses to all the tested relationships, due to an insufficient number of robustly predictive SNPs for some of the exposures. When even larger GWAS will become available, identifying more SNPs, we will be able to examine these relationships better. Third, and a more general limitation of MR, is that we cannot correct for unmeasured familial confounding, such as ‘dynastic effects’ which occur when parental genotypes have a direct effect on offspring phenotypes. This could potentially be dealt with using within-family MR studies when large enough data sets become available^53^. Fourth, the nature of our study design did not allow us to assess the role of ADHD medication status, which has previously been shown to affect substance use^5^. Fifth and final, the multiple testing burden should be considered when interpreting our findings, although this would not change our conclusions substantially, given the strong statistical evidence for the main findings.

Overall, our findings add to the current literature by allowing more robust conclusions on the causal nature of associations between ADHD and substance use. We confirm previous evidence from epidemiological studies that liability to ADHD increases the odds of initiating smoking, smoking more heavily, and finding it more difficult to quit^5^. For cannabis and alcohol use, where epidemiological studies were inconsistent, we show that liability to ADHD may increase the odds of initiating cannabis use and, tentatively, of developing alcohol dependence. This suggests that addressing ADHD symptoms early on in life may not only decrease smoking initiation and progression, but also cannabis initiation and the development of alcohol dependence. To further inform preventive efforts, future work should focus on the exact mechanisms through which causal effects of liability to ADHD are mediated. One possibility would be to perform an MR analysis for the different dimensions of ADHD (attention problems vs. impulsivity-hyperactivity) separately, if and when large enough GWAS for those phenotypes become available. Another area of interest is cognitive training. Efforts have been made to test whether training cognitive functions such as inhibitory control, which is impaired in ADHD, can decrease substance use. While for several health behaviours there is evidence that stimulus-specific inhibitory control training can be effective^54^, the literature of its efficacy on smoking is still very scarce. Our finding that smoking might causally increase ADHD risk should first be replicated and followed-up with different research methods and a wider range of measures of ADHD symptoms. Such triangulation^55^ will be essential to provide conclusive evidence on this, potentially highly impactful, finding. For the relationships where there was no indication of any causal effects – liability to ADHD and alcohol consumption and coffee use – it seems that we can, tentatively, say that the best approach for prevention would be to identify shared risk factors that are modifiable, so as to decrease risk of ADHD as well as alcohol and coffee consumption.

## Acknowledgements

MRM and HMS are members of the UK Centre for Tobacco and Alcohol Studies, a UKCRC Public Health Research: Centre of Excellence. Funding from British Heart Foundation, Cancer Research UK, Economic and Social Research Council, Medical Research Council, and National Institute for Health Research, under the auspices of the UK Clinical Research Collaboration, is gratefully acknowledged. This work was supported by the Medical Research Council Integrative Epidemiology Unit at the University of Bristol, which is supported by the Medical Research Council and the University of Bristol (grants MC_UU_12013/6 and MC_UU_12013/7). JLT is supported by a Rubicon grant from the Netherlands Organization for Scientific Research (NWO; grant number 446-16-009) as well as a Veni grant (NWO; grant number 016.Veni.195.016). KJHV and JLT are supported by the Foundation Volksbond Rotterdam. Data handling and analysis on the GenomeDK HPC facility of the ADHD GWAS was supported by NIMH (1U01MH109514-01) and Center for Genomics and Personalized Medicine (grant to ADB). ADB’s research was further supported by the Lundbeck Foundation (grant no R102-A9118 and R155-2014-1724) and by the European Community (EC) Horizon 2020 Programme (grant 667302 (CoCA)). HMS was supported by the Medical Research Council and the University of Bristol (MC_UU_00011/1, MC_UU_00011/7). Finally, we would also like to acknowledge all current members of the ADHD Working Group of the Psychiatric Genomics Consortium: Amaia Hervás, A.R. Hammerschlag, Allison Ashley-Koch, Alexandra Philipsen, Alice Charach, Ana Miranda, André Scherag, Andreas Reif, Anke Hinney, Anna Rommel, Anne Wheeler, Richard Anney, Aribert Rothenberger, Barbara Franke, Bru Cormand, Ben Neale, Christine Cornforth, Catharina Hartman, Christie Burton, Claiton Bau, Cristina Sanchez, Danielle Posthuma, Jurgen Deckert, Alysa Doyle, Eugenio Grevet, Edmund Sonuga-Barke, Elizabeth Corfield, Felecia Cerrato, Fernando Mulas, Franziska Degenhardt, Juanita Gamble, Gláucia Chiyoko Akutagava Martins, Gun Peggy Strømstad Knudsen, Hakon Hakonarson, Hans-Christoph Steinhausen, Henrik Larsson, Herber Roeyers, Peter Holmans, Jan Buitelaar, Jan Haavik, Joseph Biederman, Jennifer Crosbie, Jim McGough, Joel Gelernter, Johannes Hebebrand, Jonna Kuntsi, Joseph Sergeant, Josephine Elia, Klaus Peter Lesch, Kate Langley, Luis Rohde, Lindsey Kent, Li Yang, Maria Soler, Meg Mariano, Marieke Klein, Mark Bellgrove, Marta Ribases, Martin Steen Tesli, Joanna Martin, Miguel Casas, Michael Gill, Maria Jesús Arranz Calderón, Manuel Mattheisen, Monica Bayes, Nick Martin, Niels Peter Ole Mors, Ole Andreas Andreassen, Michael O’Donovan, Patrick Sullivan, Paul Arnold, Paul Lichtenstein, Paula Rovira, Preben Bo Mortensen, Pak Sham, Philip Asherson, Julia Pinsonneault, Patrick WL Leung, Irwin Waldman, Rachel Guerra, Josep Antoni Ramos-Quiroga, Ridha Joober, Rachel Lucier, Robert Oades, Richard Ebstein, Russell Schachar, Raymond Walters, Sarah Medland, Sarah Anthony, Sarojini Sengupta, Søren Dalsgaard, Steve Faraone, Hyo-Won Kim, Sandra Loo, Steve Nelson, Søren Dinesen, Susan Smalley, Stefan Johansson, H.C. Steinhausen, Susann Scherag, Tony Altar, Tammy Biondi, Ted Reichborn-Kjennerud, Tetyana Zayats, Anita Thapar, Tim Silk, Tinca Polderman, Tobias Banaschewski, Alexandre Todorov, Yufeng Wang, Nigel Williams, Yanil Zhang, and Ziarih Hawi.

## Authors contribution

J.L.T. carried out the analyses and drafted the manuscript. K.J.H.V and T.G.R. assisted with carrying out the analyses. All of the authors assisted with interpretation of the findings, thoroughly reviewed the content of the manuscript and approved the final version.

## References

1. Lee, S. S., Humphreys, K. L., Flory, K., Liu, R. & Glass, K. Prospective association of childhood attention-deficit/hyperactivity disorder (ADHD) and substance use and abuse/dependence: A meta-analytic review. Clin. Psychol. Rev. 31, 328–341 (2011).

2. Willcutt, E. G. The prevalence of DSM-IV attention-deficit/hyperactivity disorder: a meta-analytic review. Neurotherapeutics 9, 490–9 (2012).

3. Larsson, H., Anckarsater, H., Råstam, M., Chang, Z. & Lichtenstein, P. Childhood attention-deficit hyperactivity disorder as an extreme of a continuous trait: a quantitative genetic study of 8,500 twin pairs. J. Child Psychol. Psychiatry 53, 73–80 (2012).

4. Demontis, D. et al. Discovery of the first genome-wide significant risk loci for attention deficit/hyperactivity disorder. Nat. Genet. 51, 63–75 (2019).

5. van Amsterdam, J., van der Velde, B., Schulte, M. & van den Brink, W. Causal Factors of Increased Smoking in ADHD: A Systematic Review. Subst. Use Misuse 53, 432–445 (2018).

6. Kale, D., Stautz, K. & Cooper, A. Impulsivity related personality traits and cigarette smoking in adults: A meta-analysis using the UPPS-P model of impulsivity and reward sensitivity. Drug Alcohol Depend. 185, 149–167 (2018).

7. Mochrie, K. D., Whited, M. H., Cellucci, T., Freeman, T. & Corson, A. T. ADHD, depression, and substance abuse risk among beginning college students. J. Am. Coll. Heal. 1–5 (2018). doi:10.1080/07448481.2018.1515754

8. VanderVeen, J. D., Hershberger, A. R. & Cyders, M. A. UPPS-P model impulsivity and marijuana use behaviors in adolescents: A meta-analysis. Drug Alcohol Depend. 168, 181–190 (2016).

9. Adan, A., Forero, D. A. & Navarro, J. F. Personality Traits Related to Binge Drinking: A Systematic Review. Front. Psychiatry 8, 134 (2017).

10. Marmorstein, N. R. Energy Drink and Coffee Consumption and Psychopathology Symptoms Among Early Adolescents: Cross-Sectional and Longitudinal Associations. J. Caffeine Res. 6, 64–72 (2016).

11. Dosh, T. et al. A Comparison of the Associations of Caffeine and Cigarette Use With Depressive and ADHD Symptoms in a Sample of Young Adult Smokers. J. Addict. Med. 4, 50–52 (2010).

12. Konstenius, M. et al. Childhood trauma exposure in substance use disorder patients with and without ADHD. Addict. Behav. 65, 118–124 (2017).

13. Green, J. G. et al. Childhood Adversities and Adult Psychiatric Disorders in the National Comorbidity Survey Replication I. Arch. Gen. Psychiatry 67, 113 (2010).

14. Elkins, I. J. et al. Associations between childhood ADHD, gender, and adolescent alcohol and marijuana involvement: A causally informative design. Drug Alcohol Depend. 184, 33–41 (2018).

15. Skoglund, C., Chen, Q., Franck, J., Lichtenstein, P. & Larsson, H. Attention-Deficit/Hyperactivity Disorder and Risk for Substance Use Disorders in Relatives. Biol. Psychiatry 77, 880–886 (2015).

16. Pasman, J. A. et al. GWAS of lifetime cannabis use reveals new risk loci, genetic overlap with psychiatric traits, and a causal influence of schizophrenia. Nat. Neurosci. 21, 1161–1170 (2018).

17. Liu, M. et al. Association studies of up to 1.2 million individuals yield new insights into the genetic etiology of tobacco and alcohol use. Nat. Genet. 1 (2019). doi:10.1038/s41588-018-0307-5

18. Cornelis, M. C. et al. Genome-wide meta-analysis identifies six novel loci associated with habitual coffee consumption. Mol. Psychiatry 20, 647–656 (2015).

19. Sanchez-Roige, S. et al. Genome-wide association study meta-analysis of the Alcohol Use Disorder Identification Test (AUDIT) in two population-based cohorts (N=141,932). bioRxiv 275917 (2018). doi:10.1101/275917

20. Rubia, K. Cognitive Neuroscience of Attention Deficit Hyperactivity Disorder (ADHD) and Its Clinical Translation. Front. Hum. Neurosci. 12, 100 (2018).

21. Sampedro-Piquero, P. et al. Neuroplastic and cognitive impairment in substance use disorders: a therapeutic potential of cognitive stimulation. Neurosci. Biobehav. Rev. (2018). doi:10.1016/j.neubiorev.2018.11.015

22. Arcos-Burgos, M., Vélez, J. I., Solomon, B. D. & Muenke, M. A common genetic network underlies substance use disorders and disruptive or externalizing disorders. Hum. Genet. 131, 917–29 (2012).

23. Norén Selinus, E. et al. Subthreshold and threshold attention deficit hyperactivity disorder symptoms in childhood: psychosocial outcomes in adolescence in boys and girls. Acta Psychiatr. Scand. 134, 533–545 (2016).

24. Treur, JL, Willemsen, G, Bartels, M, Geels, LM, van Beek, JHDA, Huppertz, C, van Beijsterveldt CEM, Boomsma, DI, Vink, J. Smoking During Adolescence as a Risk Factor for Attention Problems. Biol. Psychiatry 78, 656–663 (2015).

25. Pardini, D. et al. Unfazed or Dazed and Confused: Does Early Adolescent Marijuana Use Cause Sustained Impairments in Attention and Academic Functioning? J. Abnorm. Child Psychol. 43, 1203–1217 (2015).

26. Kelly, C. et al. Distinct effects of childhood ADHD and cannabis use on brain functional architecture in young adults. NeuroImage Clin. 13, 188–200 (2017).

27. Louth, E. L., Bignell, W., Taylor, C. L. & Bailey, C. D. C. Developmental Ethanol Exposure Leads to Long-Term Deficits in Attention and Its Underlying Prefrontal Circuitry. eNeuro 3, (2016).

28. Lawlor, D. A., Harbord, R. M., Sterne, J. A. C., Timpson, N. & Davey Smith, G. Mendelian randomization: Using genes as instruments for making causal inferences in epidemiology. Stat. Med. 27, 1133–1163 (2008).

29. Davies, N. M., Holmes, M. V & Davey Smith, G. Reading Mendelian randomisation studies: a guide, glossary, and checklist for clinicians. BMJ 362, k601 (2018).

30. Chao, M., Li, X. & McGue, M. The Causal Role of Alcohol Use in Adolescent Externalizing and Internalizing Problems: A Mendelian Randomization Study. Alcohol. Clin. Exp. Res. 41, 1953–1960 (2017).

31. Fluharty, M. E., Sallis, H. & Munafò, M. R. Investigating possible causal effects of externalizing behaviors on tobacco initiation: A Mendelian randomization analysis. Drug Alcohol Depend. 191, 338–342 (2018).

32. Sallis, H. M., Smith, G. D. & Munafo, M. R. Cigarette smoking and personality: Investigating causality using Mendelian randomization. Psychol Med 25, 1–9 (2018).

33. Soler Artigas, M. et al. Attention-deficit/hyperactivity disorder and lifetime cannabis use: genetic overlap and causality. Mol. Psychiatry 1 (2019). doi:10.1038/s41380-018-0339-3

34. Pingault, J.-B. et al. Using genetic data to strengthen causal inference in observational research. Nat. Rev. Genet. 19, 566–580 (2018).

35. Wootton, R. E. et al. Causal effects of lifetime smoking on risk for depression and schizophrenia: Evidence from a Mendelian randomisation study. bioRxiv 381301 (2018). doi:10.1101/381301

36. Walters, R. K. et al. Transancestral GWAS of alcohol dependence reveals common genetic underpinnings with psychiatric disorders. Nat. Neurosci. 21, 1656–1669 (2018).

37. Hemani, G. et al. The MR-Base platform supports systematic causal inference across the human phenome. Elife 7, (2018).

38. Burgess, S. & Davey Smith, G. How humans can contribute to Mendelian randomization analyses. Int. J. Epidemiol. (2019). doi:10.1093/ije/dyz152

39. Bowden, J., Davey Smith, G., Haycock, P. C. & Burgess, S. Consistent Estimation in Mendelian Randomization with Some Invalid Instruments Using a Weighted Median Estimator. Genet. Epidemiol. 40, 304–314 (2016).

40. Hartwig, F. P., Smith, G. D. & Bowden, J. Robust inference in two-sample Mendelian randomisation via the zero modal pleiotropy assumption. Int. J. Epidemiol. 46, 1985–1998 (2017).

41. Bowden, J., Davey Smith, G. & Burgess, S. Mendelian randomization with invalid instruments: effect estimation and bias detection through Egger regression. Int. J. Epidemiol. 44, 512–525 (2015).

42. Bowden, J. et al. Assessing the suitability of summary data for two-sample Mendelian randomization analyses using MR-Egger regression: the role of the I2 statistic. Int. J. Epidemiol. 45, dyw220 (2016).

43. Zhu, Z. et al. Causal associations between risk factors and common diseases inferred from GWAS summary data. Nat. Commun. 9, 224 (2018).

44. Hemani, G., Tilling, K. & Davey Smith, G. Orienting The Causal Relationship Between Imprecisely Measured Traits Using Genetic Instruments. PLOS Genet. 13, e1007081 (2017).

45. O’Connor, L. J. & Price, A. L. Distinguishing genetic correlation from causation across 52 diseases and complex traits. Nat. Genet. 50, 1728–1734 (2018).

46. Bowden, J., Hemani, G. & Davey Smith, G. Invited Commentary: Detecting Individual and Global Horizontal Pleiotropy in Mendelian Randomization—A Job for the Humble Heterogeneity Statistic? Am. J. Epidemiol. 187, 2681–2685 (2018).

47. Marees AT, Smit DJA, Ong J-S, MacGregor S, An J, Denys D, Vorspan F, van den Brink W, D. E. Potential influence of socio-economic status on genetic correlations between alcohol consumption measures and mental health. Psychol Med 15, 1–15 (2019).

48. Middeldorp, C. M. et al. A Genome-Wide Association Meta-Analysis of Attention-Deficit/Hyperactivity Disorder Symptoms in Population-Based Pediatric Cohorts. J. Am. Acad. Child Adolesc. Psychiatry 55, 896–905.e6 (2016).

49. Egan, T. E., Dawson, A. E. & Wymbs, B. T. Substance Use in Undergraduate Students With Histories of Attention-Deficit/Hyperactivity Disorder (ADHD): The Role of Impulsivity. Subst. Use Misuse 52, 1375–1386 (2017).

50. Taylor, G. M. J. & Munafò, M. R. Does smoking cause poor mental health? The Lancet Psychiatry 6, 2–3 (2019).

51. Counotte, D. S., Smit, A. B. & Spijker, S. The Yin and Yang of Nicotine: Harmful during Development, Beneficial in Adult Patient Populations. Front. Pharmacol. 3, 180 (2012).

52. Cooper, M. et al. Investigating late-onset ADHD: a population cohort investigation. J. Child Psychol. Psychiatry 59, 1105–1113 (2018).

53. Brumpton, B. et al. Within-family studies for Mendelian randomization: avoiding dynastic, assortative mating, and population stratification biases. bioRxiv 602516 (2019). doi:10.1101/602516

54. Allom, V., Mullan, B. & Hagger, M. Does inhibitory control training improve health behaviour? A meta-analysis. Health Psychol. Rev. 10, 168–186 (2016).

55. Lawlor, D. A., Tilling, K. & Davey Smith, G. Triangulation in aetiological epidemiology. Int. J. Epidemiol. 45, dyw314 (2017).

