## Supplemental material for "Investigating causality between liability to ADHD and substance use, and liability to substance use and ADHD risk, using Mendelian randomization"

| | |
|---|---|
| Page 2 | Supplementary Table 1. F statistic, indicating instrument strength |
| Page 3 | Supplementary Table 2. MR-Egger intercept, indicating horizontal pleiotropy |
| Page 4 | Supplementary Table 3. Cochran's heterogeneity statistic |
| Page 5 | Supplementary Table 4. $I^2$ statistic |
| Page 6 | Supplementary Table 5. Mendelian randomization analyses from liability to ADHD to alcohol intake frequency, in UK Biobank |
| Page 7 | Supplementary Table 6. Mendelian randomization analyses after Steiger filtering |
| Page 8 | Supplementary Table 7. Mendelian randomization analyses from liability to smoking initiation to *children* (<13 years) ADHD symptoms |
| Page 9 | Supplementary Table 8. Results of LCV analyses |
| Page 10-25 | Supplementary Figure 1 – Figure 15. Leave-one-out IVW Mendelian randomization analyses |

**Supplementary Table 1.** F statistic, indicating instrument strength, for Mendelian randomization analyses from liability to ADHD to substance use risk, and from liability to substance use to ADHD risk.

| Exposure | Outcome | *n* SNPs | Mean F |
|---|---|---|---|
| ADHD | Smoking initiation | 10 | 33.65 |
| ADHD | Cigarettes / day | 10 | 33.65 |
| ADHD | Smoking cessation | 11 | 33.66 |
| ADHD | Lifetime smoking | 10 | 34.08 |
| ADHD | Alcohol drinks / week | 10 | 33.65 |
| ADHD | Alcohol problems | 10 | 33.65 |
| ADHD | Alcohol dependence | 12 | 33.66 |
| ADHD | Cannabis initiation | 10 | 34.08 |
| ADHD | Cups of coffee / day | 9 | 33.31 |
| Smoking initiation | ADHD | 346 | 45.15 |
| Alcohol drinks / week | ADHD | 90 | 64.08 |
| Alcohol problems | ADHD | 7 | 48.03 |
| Alcohol dependence | ADHD | 9 | 29.73 |
| Cannabis initiation | ADHD | 5 | 40.55 |
| Cups of coffee / day | ADHD | 4 | 134.69 |

*n* SNPs = number of SNPs included in the genetic instrument. The F-statistic quantifies instrumental bias, where F>10 indicates the instrument is sufficiently strong

**Supplementary Table 2.** MR-Egger intercept, indicating horizontal pleiotropy, for Mendelian randomization analyses from liability to ADHD to substance use risk, and from liability to substance use to ADHD risk.

| Exposure | Outcome | *n* SNPs | MR-Egger intercept | | |
|---|---|---|---|---|---|
| | | | beta | 95% CI | *p* |
| ADHD | Smoking initiation | 10 | 0.01 | -0.01 to 0.02 | 0.412 |
| ADHD | Cigarettes / day | 10 | 0.01 | 0.00 to 0.02 | 0.068 |
| ADHD | Smoking cessation | 11 | 0.01 | 0.00 to 0.01 | 0.089 |
| ADHD | Lifetime smoking | 10 | *n.a.* | *n.a.* | *n.a.* |
| ADHD | Alcoholic drinks / week | 10 | -0.01 | -0.03 to 0.01 | 0.468 |
| ADHD | Alcohol problems | 10 | *n.a.* | *n.a.* | *n.a.* |
| ADHD | Alcohol dependence | 12 | 0.02 | -0.02 to 0.06 | 0.226 |
| ADHD | Cannabis initiation | 10 | *n.a.* | *n.a.* | *n.a.* |
| Smoking initiation* | ADHD | 346 | 0.01 | -0.01 to 0.03 | 0.375 |
| Alcohol drinks / week | ADHD | 90 | 0.01 | -0.01 to 0.03 | 0.372 |
| Alcohol problems | ADHD | 7 | *n.a.* | *n.a.* | *n.a.* |
| Alcohol dependence | ADHD | 9 | *n.a.* | *n.a.* | *n.a.* |
| Cannabis initiation | ADHD | 5 | *n.a.* | *n.a.* | *n.a.* |
| Cups of coffee / day | ADHD | 4 | *n.a.* | *n.a.* | *n.a.* |

*n* SNPs = number of SNPs included in the genetic instrument**;** SE = standard error of the intercept. The dichotomous variables smoking initiation and smoking cessation were rescaled in the original GWAS such that its unit is a standard deviation increase in prevalence[32]. For MR-Egger; when $I^2$ was 0.6-0.9, a SIMEX correction was applied, while estimates were not reported at all when $I^2$ was <0.6. *n.a:* the number of SNPs available for the analysis was too low or $I^2$ was <0.6.

**Supplementary Table 3.** Cochran's heterogeneity statistic for Inverse Variance Weighted (IVW) Mendelian randomization analyses from liability to ADHD to substance use risk, and from liability to substance use to ADHD risk.

| Exposure | Outcome | *n* SNPs | Q | *p* |
|---|---|---|---|---|
| ADHD | Smoking initiation | 10 | 34.44 | 7.5e-05 |
| ADHD | Cigarettes / day | 10 | 11.33 | 0.254 |
| ADHD | Smoking cessation | 11 | 11.76 | 0.302 |
| ADHD | Lifetime smoking | 10 | 46.88 | 4.1e-07 |
| ADHD | Alcohol drinks / week | 10 | 39.24 | 1.0e-05 |
| ADHD | Alcohol problems | 10 | 18.39 | 0.031 |
| ADHD | Alcohol dependence | 12 | 10.41 | 0.494 |
| ADHD | Cannabis initiation | 10 | 15.90 | 0.069 |
| ADHD | Cups of coffee / day | 9 | 6.73 | 0.566 |
| Smoking initiation | ADHD | 346 | 373.84 | 0.137 |
| Alcohol drinks / week | ADHD | 90 | 115.87 | 0.029 |
| Alcohol problems | ADHD | 7 | 7.47 | 0.279 |
| Alcohol dependence | ADHD | 9 | 9.86 | 0.275 |
| Cannabis initiation | ADHD | 5 | 5.82 | 0.213 |
| Cups of coffee / day | ADHD | 4 | 6.65 | 0.084 |

*n* SNPs = number of SNPs included in the genetic instrument.

**Supplementary Table 4.** $I^2$ statistic for Mendelian randomization analyses from liability to ADHD to substance use risk, and from liability to substance use to ADHD risk.

| Exposure | Outcome | *n* SNPs | $I^2$ |
|---|---|---|---|
| ADHD | Smoking initiation | 10 | 0.93 |
| ADHD | Cigarettes / day | 10 | 0.81 |
| ADHD | Smoking cessation | 11 | 0.94 |
| ADHD | Lifetime smoking | 10 | 0.39 |
| ADHD | Alcohol drinks / week | 10 | 0.80 |
| ADHD | Alcohol problems | 10 | 0.28 |
| ADHD | Alcohol dependence | | |
| ADHD | Cannabis initiation | 10 | 0.38 |
| ADHD | Cups of coffee / day | 9 | *n.a.* |
| Smoking initiation | ADHD | 346 | 0.60 |
| Alcohol drinks / week | ADHD | 90 | 0.95 |
| Alcohol problems | ADHD | 7 | *n.a.* |
| Alcohol dependence | ADHD | 9 | *n.a.* |
| Cannabis initiation | ADHD | 5 | 0.76 |
| Cups of coffee / day | ADHD | 4 | *n.a.* |

*n* SNPs = number of SNPs included in the genetic instrument. $I^2$ = statistic that quantifies heterogeneity between the genetic variants in an instrument and indicates whether the 'NO Measurement Error' (NOME) assumption has been violated. An $I^2$ value below 0.9 means that there is a considerable risk of bias but it may still be corrected for by applying MR-Egger simulation extrapolation (SIMEX). An $I^2$ value below 0.6 means that the results of MR-Egger (even with SIMEX correction) are unreliable.

**Supplementary Table 5.** Mendelian randomization analysis from liability to ADHD to alcohol intake frequency, in UK Biobank (*n* = 336,965).

| Exposure | Outcome | | IVW | | | Weighted median | | | Weighted mode | | | MR-Egger | | | GSMR | | | |
|---|---|---|---|---|---|---|---|---|---|---|---|---|---|---|---|---|---|---|
| | | *n* SNPs | beta | 95% CI | *p* | beta | 95% CI | *p* | beta | 95% CI | *p* | beta | 95% CI | *p* | *n* SNPs | beta | 95% CI | *p* |
| ADHD | Alcohol intake frequency | 8 | 0.22 | 0.04 to 0.40 | 0.014 | 0.20 | 0.08 to 0.32 | 3.4e-4 | 0.05 | -0.01 to 0.11 | 0.118 | *n.a.* | *n.a.* | *n.a.* | 8 | *n.a.* | *n.a.* | *n.a.* |

For MR-Egger; when $I^2$ was 0.6-0.9, a SIMEX correction was applied, while estimates were not reported at all when $I^2$ was <0.6. *n.a:* the number of SNPs available for the analysis was too low, or, in the case of MR-Egger, $I^2$ was <0.6.

**Supplementary Table 6.** Results of the Mendelian randomization analyses from liability to ADHD to substance use risk and from liability to substance use to ADHD risk, including Inverse Variance Weighted (IVW) estimates and four sensitivity analyses: weighted median, weighted mode, MR-Egger regression and GSMR (generalized summary-data-based Mendelian randomization) – ***after Steiger filtering***

| Exposure | Outcome | *n* | IVW | | | | Weighted median | | | | Weighted mode | | | | MR-Egger | | | | *n** | GSMR | | | |
|---|---|---|---|---|---|---|---|---|---|---|---|---|---|---|---|---|---|---|---|---|---|---|---|
| | | SNPs | beta | OR | 95% CI | *p* | beta | OR | 95% CI | *p* | beta | OR | 95% CI | *p* | beta | OR | 95% CI | *p* | SNPs | beta | OR | 95% CI | *p* |
| ADHD | Smoking initiation | 9 | 0.07 | | 0.05 to 0.09 | 1.9e-6 | 0.05 | | 0.03 to 0.07 | 5.4e-5 | 0.05 | | 0.01 to 0.09 | 0.021 | *n.a.* | | *n.a.* | *n.a.* | | *n.a.* | | *n.a.* | *n.a.* |
| ADHD | Cigarettes / day | 10 | | | | | | | | | | | *unchanged* | | | | | | | | | | |
| ADHD | Smoking cessation | 11 | | | | | | | | | | | *unchanged* | | | | | | | | | | |
| ADHD | Lifetime smoking | 10 | | | | | | | | | | | *unchanged* | | | | | | | | | | |
| ADHD | Alcohol drinks / week | 10 | | | | | | | | | | | *unchanged* | | | | | | | | | | |
| ADHD | Alcohol problems | 10 | | | | | | | | | | | *unchanged* | | | | | | | | | | |
| ADHD | Alcohol dependence | 12 | | | | | | | | | | | *unchanged* | | | | | | | | | | |
| ADHD | Cannabis initiation | 10 | | | | | | | | | | | *unchanged* | | | | | | | | | | |
| ADHD | Cups of coffee / day | 9 | | | | | | | | | | | *unchanged* | | | | | | | | | | |
| Smoking initiation | ADHD | 265 | 0.79 | 2.21 | 1.84 to 2.64 | 3.5e-17 | 0.89 | 2.43 | 1.86 to 3.19 | 8.4e-11 | 1.07 | 2.92 | 1.16 to 7.32 | 0.022 | 1.34 | 3.82 | 2.34 to 6.23 | 1.7e-7 | 255 | 0.77 | 2.16 | 1.77 to 2.64 | 2.2e-15 |
| Alcohol drinks / week | ADHD | 49 | -0.06 | 0.94 | 0.50 to 1.77 | 0.837 | -0.06 | 0.94 | 0.41 to 2.18 | 0.886 | -0.49 | 0.61 | 0.22 to 1.70 | 0.344 | -0.21 | 0.81 | -1.29 to 0.87 | 0.711 | 43 | -0.02 | 0.98 | 0.49 to 1.95 | 0.954 |
| Alcohol problems | ADHD | 4 | 0.39 | 1.47 | 0.09 to 23.34 | 0.783 | 0.08 | 1.08 | 0.03 to 26.84 | 0.961 | -0.37 | 0.69 | 0.01 to 43.38 | 0.873 | *n.a.* | | *n.a.* | *n.a.* | | *n.a.* | | *n.a.* | *n.a.* |
| Alcohol dependence | ADHD | 9 | | | | | | | | | | | *unchanged* | | | | | | | | | | |
| Cannabis initiation | ADHD | 5 | | | | | | | | | | | *unchanged* | | | | | | | | | | |
| Cups of coffee / day | ADHD | 4 | | | | | | | | | | | *unchanged* | | | | | | | | | | |

*n* SNPs = number of SNPs included in the genetic instrument**.** SE = standard error of the beta. Note that the dichotomous variables smoking initiation and smoking cessation were rescaled in the original GWAS such that its unit is a standard deviation increase in prevalence[28]. For MR-Egger; when $I^2$ was 0.6-0.9, a SIMEX correction was applied, while estimates were not reported at all when $I^2$ was <0.6. *n.a:* the number of SNPs available for the analysis was too low, or, in the case of MR-Egger, $I^2$ was <0.6. ***Number of SNPs left after the HEIDI filtering step which is part of GSMR.

**Supplementary table 7.** Results of the Mendelian randomization analyses using summary level data from liability to smoking initiation to *children* (<13 years) ADHD symptoms including IVW estimates and four sensitivity analyses: weighted median, weighted mode, MR-Egger, and GSMR (generalized summary-level-data based Mendelian randomization).

| Exposure | Outcome | IVW | | | | Weighted median | | | Weighted mode | | | MR-Egger | | | GSMR | | | |
|---|---|---|---|---|---|---|---|---|---|---|---|---|---|---|---|---|---|---|
| | | *n* SNPs | beta | 95% CI | *p* | beta | 95% CI | *p* | beta | 95% CI | *p* | beta | 95% CI | *p* | *n* SNPs* | beta | 95% CI | *p* |
| Smoking initiation | ADHD | 296 | 0.28 | 0.17 to 0.39 | 3.5e-07 | 0.23 | 0.08 to 0.39 | 0.004 | 0.39 | -0.03 to 0.81 | 0.075 | *n.a.* | *n.a.* | *n.a.* | 289 | 0.25 | 0.15 to 0.35 | 1.4e-06 |

*n* SNPs = number of SNPs included in the genetic instrument**.** SE = standard error of the beta. Note that the dichotomous variables smoking initiation and smoking cessation were rescaled in the original GWAS such that its unit is a standard deviation increase in prevalence[28]. For MR-Egger; when $I^2$ was 0.6-0.9, a SIMEX correction was applied, while estimates were not reported at all when $I^2$ was <0.6. *n.a:* the number of SNPs available for the analysis was too low, or, in the case of MR-Egger, $I^2$ was <0.6. *****Number of SNPs left after the HEIDI filtering step which is part of GSMR.

**Supplementary Table 8.** Results of the LCV (Latent Causal Variable) model

| Trait 1 | Trait 2 | *n* SNPs | GCP | p-value GCP | p-value to reject H0 that GCP=1 | p-value to reject H0 that GCP=-1 |
|---|---|---|---|---|---|---|
| ADHD | Smoking initiation | 1,177,032 | -0.82 | 6.6e-08 | 7.4e-06 | 0.951 |
| ADHD | Alcohol drinks / week | 1,184,013 | 0.19 | 0.685 | 0.695 | 0.924 |
| ADHD | Alcohol problems | 1,190,000 | -0.07 | 0.585 | 0.242 | 0.479 |
| ADHD | Alcohol dependence | 1,199,049 | -0.57 | 0.004 | 1 | 0.687 |
| ADHD | Cannabis initiation | 1,195,821 | 0.01 | 0.730 | 0.529 | 0.384 |
| ADHD | Coffee / day | 1,072,163 | -0.07 | 0.317 | 0.821 | 0.335 |

The LCV model uses SNPs from across the whole genome (no p-value threshold is imposed) and it does *not* indicate which of the two traits is the exposure and which is the outcome variable. A latent causal variable (LCV) Is modelled which mediates the genetic correlation between trait 1 and trait 2. If trait 1 is strongly genetically correlated with the latent causal variable, this means that trait 1 is genetically causal for trait 2; if trait 2 is genetically correlated with the latent causal variable, this means that trait 2 is genetically causal for trait 1. The GCP (Genetic Causality Proportion) statistic reflects which of the two traits is genetically correlated with the latent causal variable. If GCP = 1, this implies that trait 1 is genetically causal for trait 2, if GCP = -1, this implies that trait 2 is genetically causal for trait 1. Note that ADHD here is based on the complete GWAS summary statistics, including children, adolescents and adults.

**Supplementary Figure 1.** Leave-one-out analyses depicting the results of Inverse Variance Weighted (IVW) Mendelian randomization analyses from liability to ADHD to smoking initiation risk after excluding each of the genetic variants from the analysis, one at the time. This gives an indication of whether or not the overall effect is driven by one particular genetic variant.

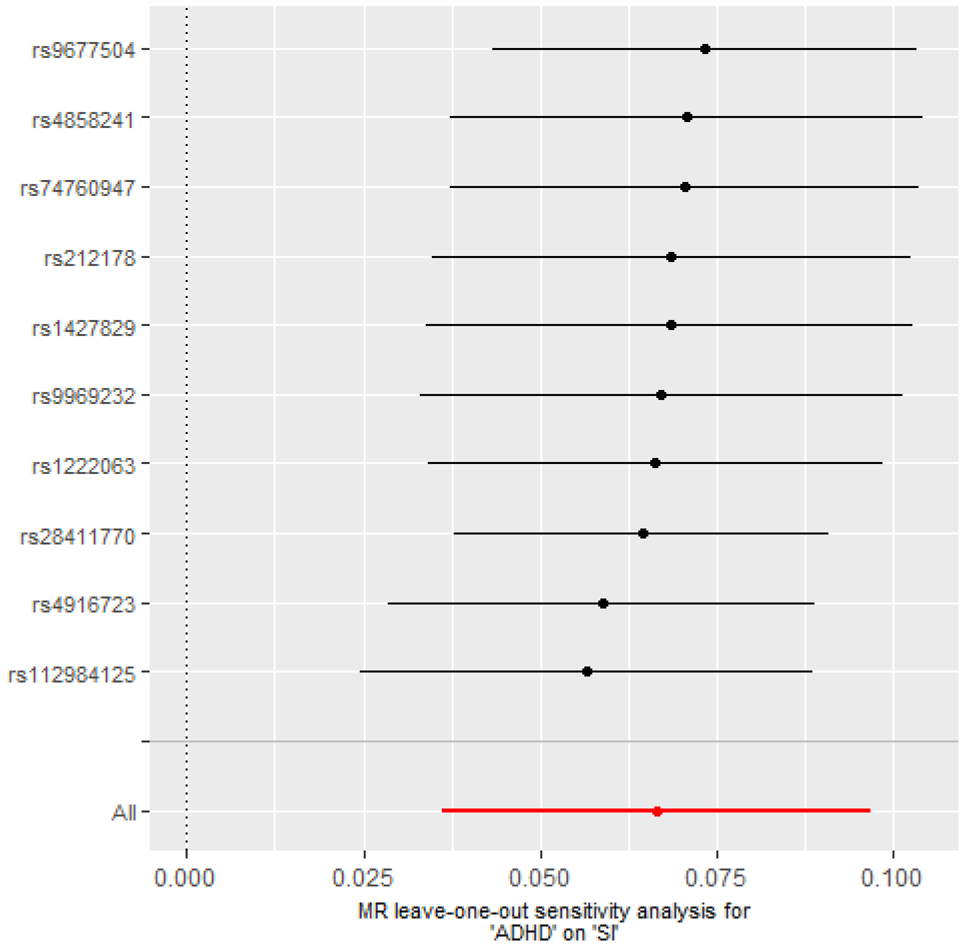

**Supplementary Figure 2.** Leave-one-out analyses depicting the results of Inverse Variance Weighted (IVW) Mendelian randomization analyses from liability to ADHD to cigarettes smoked per day after excluding each of the genetic variants from the analysis, one at the time. This gives an indication of whether or not the overall effect is driven by one particular genetic variant.

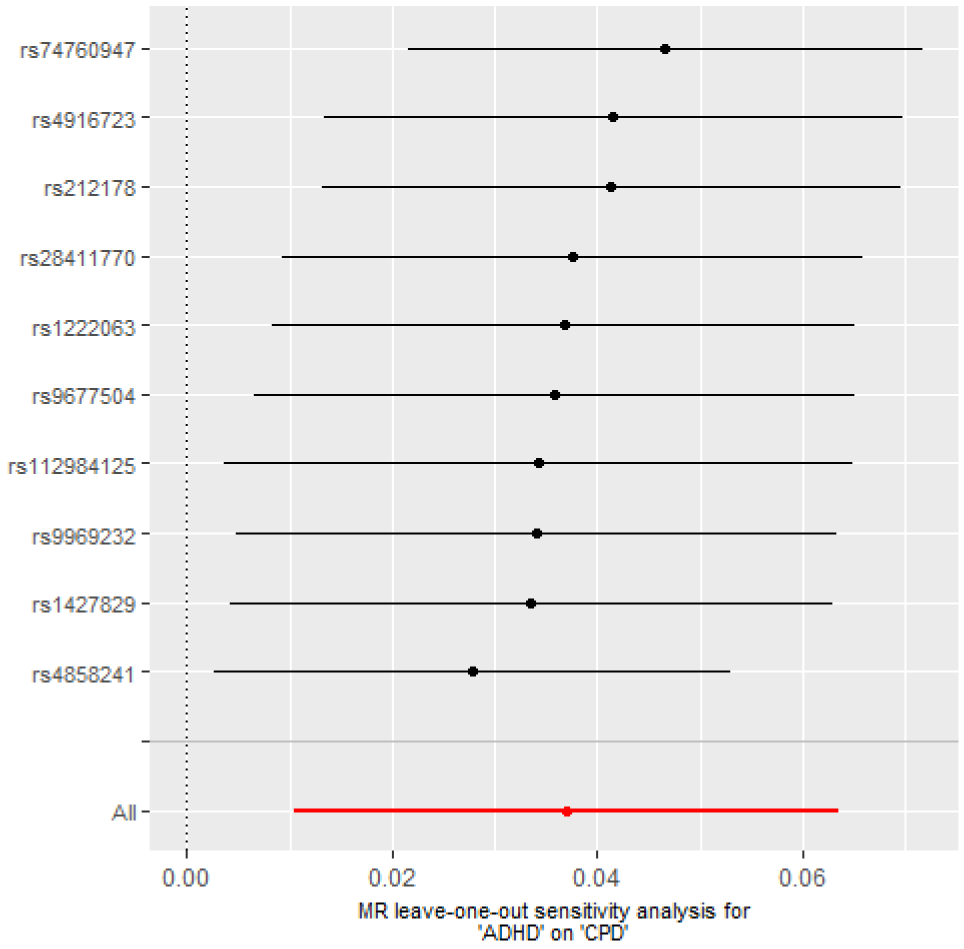

**Supplementary Figure 3.** Leave-one-out analyses depicting the results of Inverse Variance Weighted (IVW) Mendelian randomization analyses from liability to ADHD to smoking cessation risk after excluding each of the genetic variants from the analysis, one at the time. This gives an indication of whether or not the overall effect is driven by one particular genetic variant.

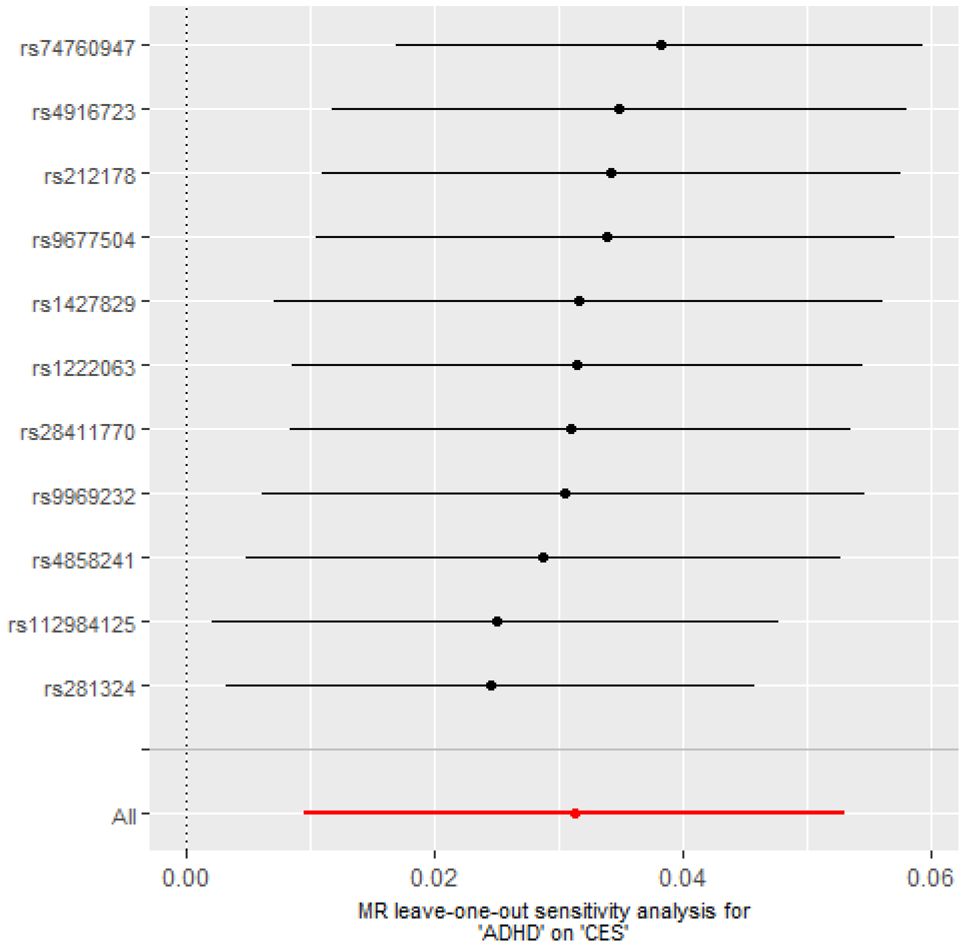

**Supplementary Figure 4.** Leave-one-out analyses depicting the results of Inverse Variance Weighted (IVW) Mendelian randomization analyses from liability to ADHD to lifetime smoking after excluding each of the genetic variants from the analysis, one at the time. This gives an indication of whether or not the overall effect is driven by one particular genetic variant.

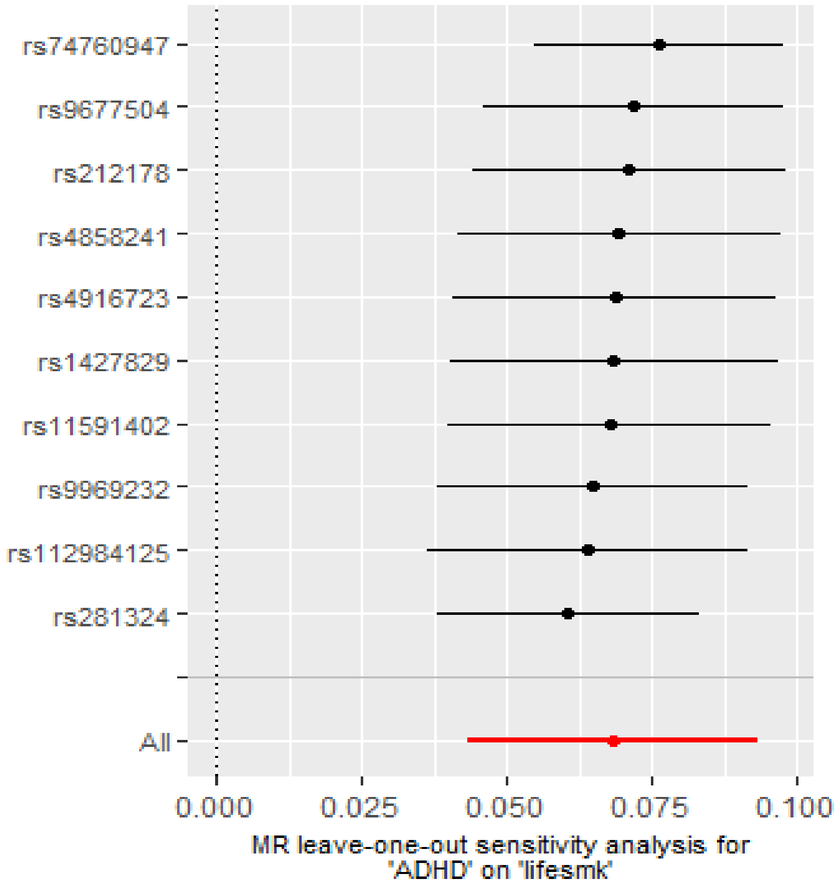

**Supplementary Figure 5.** Leave-one-out analyses depicting the results of Inverse Variance Weighted (IVW) Mendelian randomization analyses from liability to ADHD to alcohol drinks per week after excluding each of the genetic variants from the analysis, one at the time. This gives an indication of whether or not the overall effect is driven by one particular genetic variant.

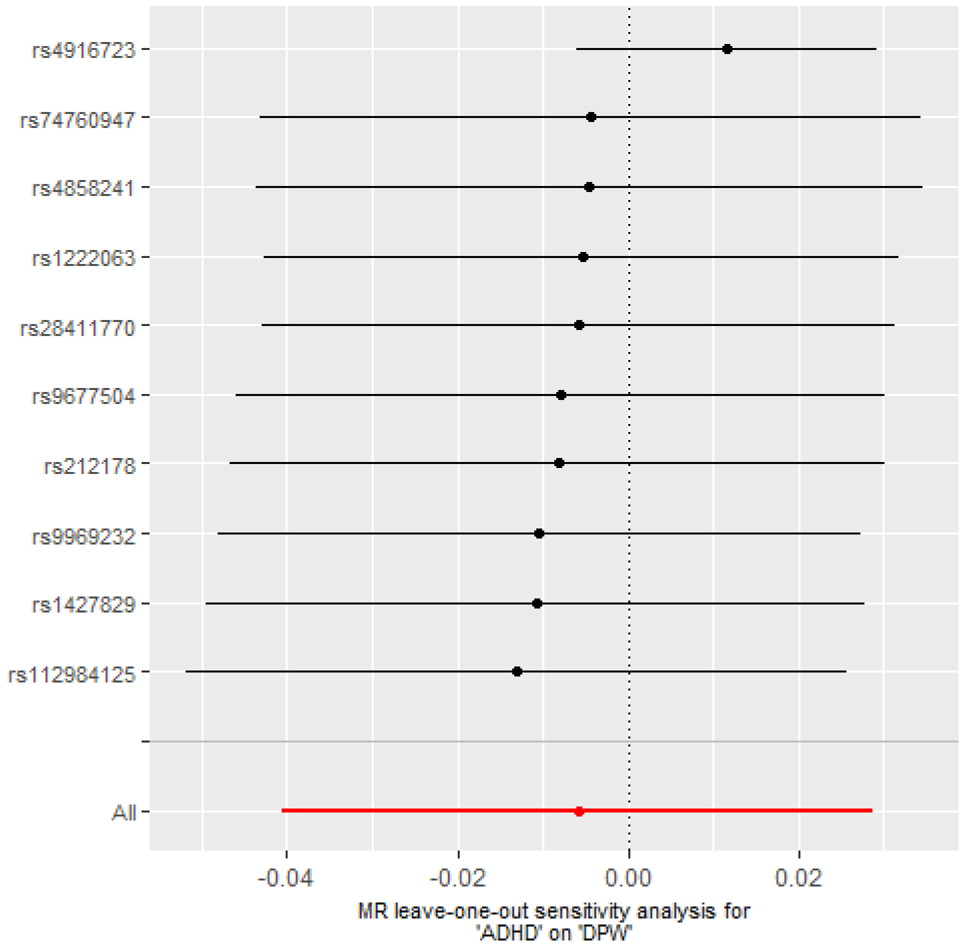

**Supplementary Figure 6.** Leave-one-out analyses depicting the results of Inverse Variance Weighted (IVW) Mendelian randomization analyses from liability to ADHD to alcohol problems risk after excluding each of the genetic variants from the analysis, one at the time. This gives an indication of whether or not the overall effect is driven by one particular genetic variant.

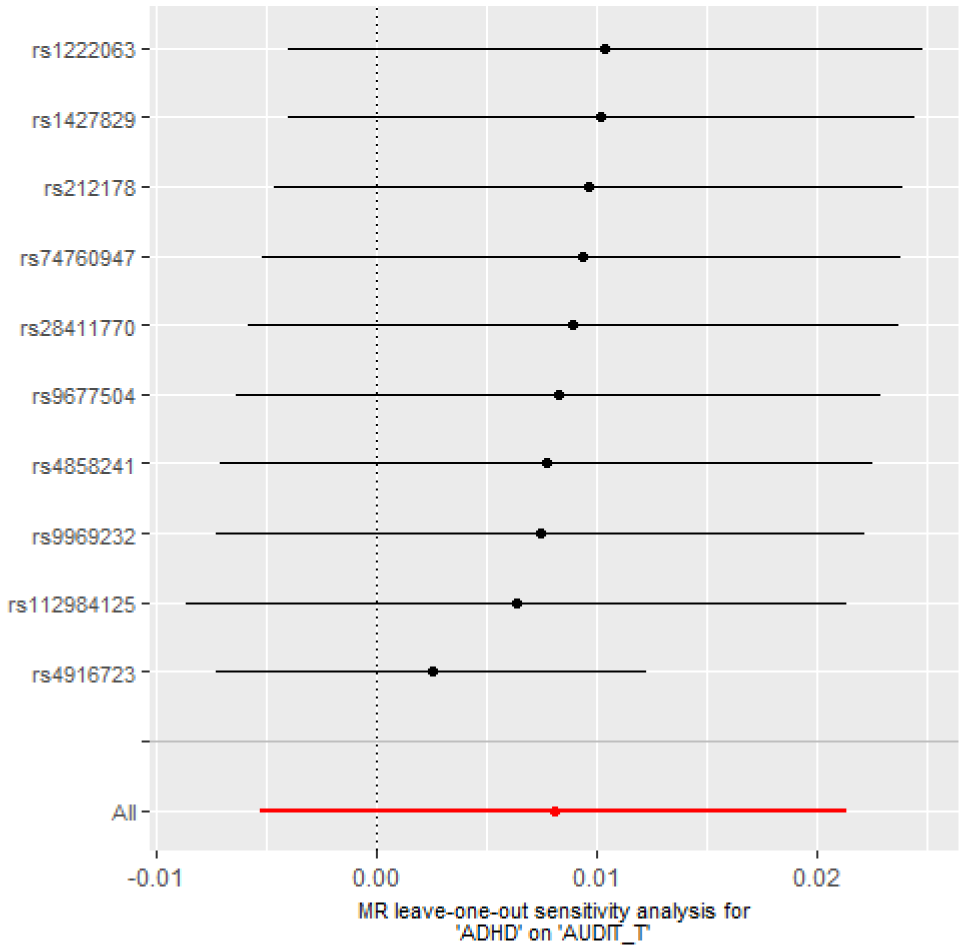

**Supplementary Figure 7.** Leave-one-out analyses depicting the results of Inverse Variance Weighted (IVW) Mendelian randomization analyses from liability to ADHD to alcohol dependence risk after excluding each of the genetic variants from the analysis, one at the time. This gives an indication of whether or not the overall effect is driven by one particular genetic variant.

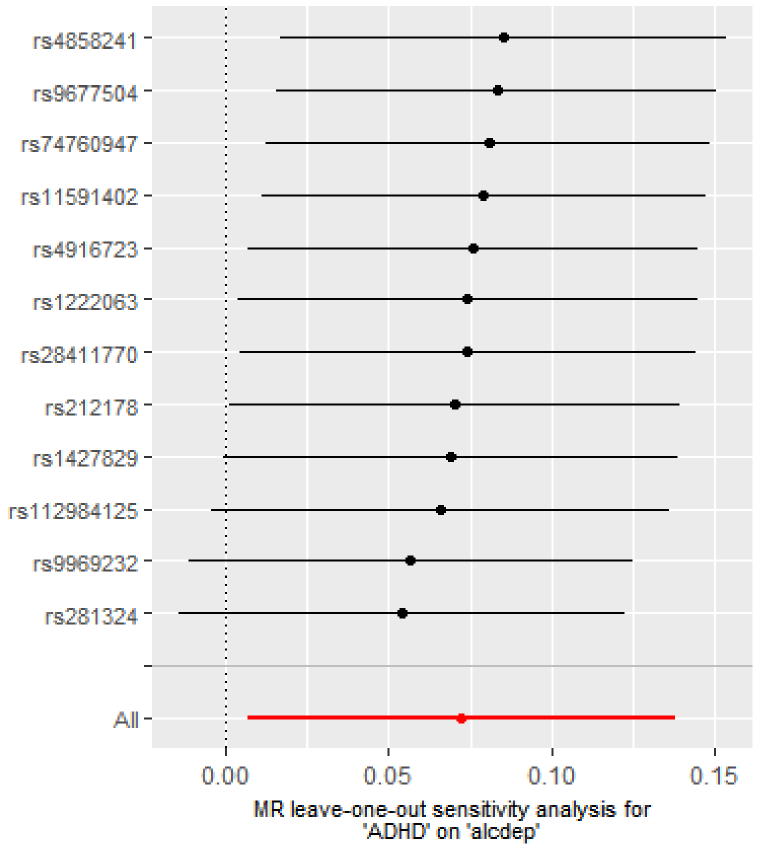

**Supplementary Figure 8.** Leave-one-out analyses depicting the results of Inverse Variance Weighted (IVW) Mendelian randomization analyses from liability to ADHD to cannabis initiation risk after excluding each of the genetic variants from the analysis, one at the time. This gives an indication of whether or not the overall effect is driven by one particular genetic variant.

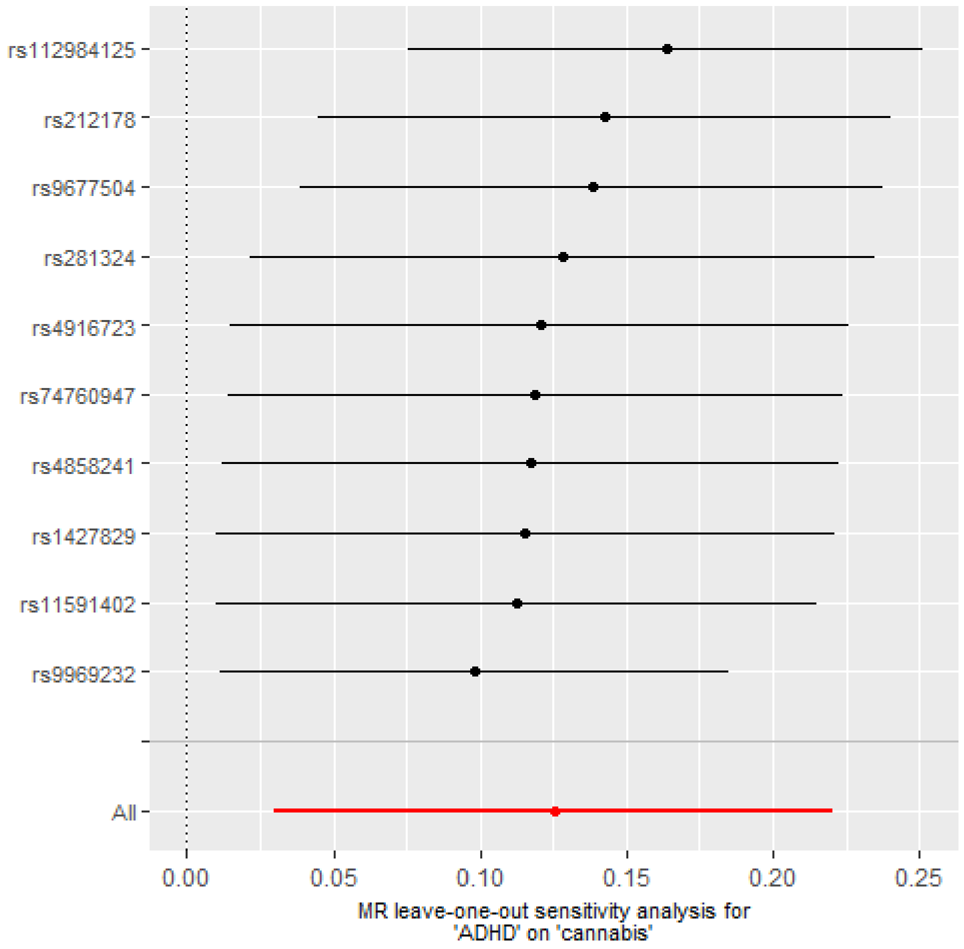

**Supplementary Figure 9.** Leave-one-out analyses depicting the results of Inverse Variance Weighted (IVW) Mendelian randomization analyses from liability to ADHD to cups of coffee per day after excluding each of the genetic variants from the analysis, one at the time. This gives an indication of whether or not the overall effect is driven by one particular genetic variant.

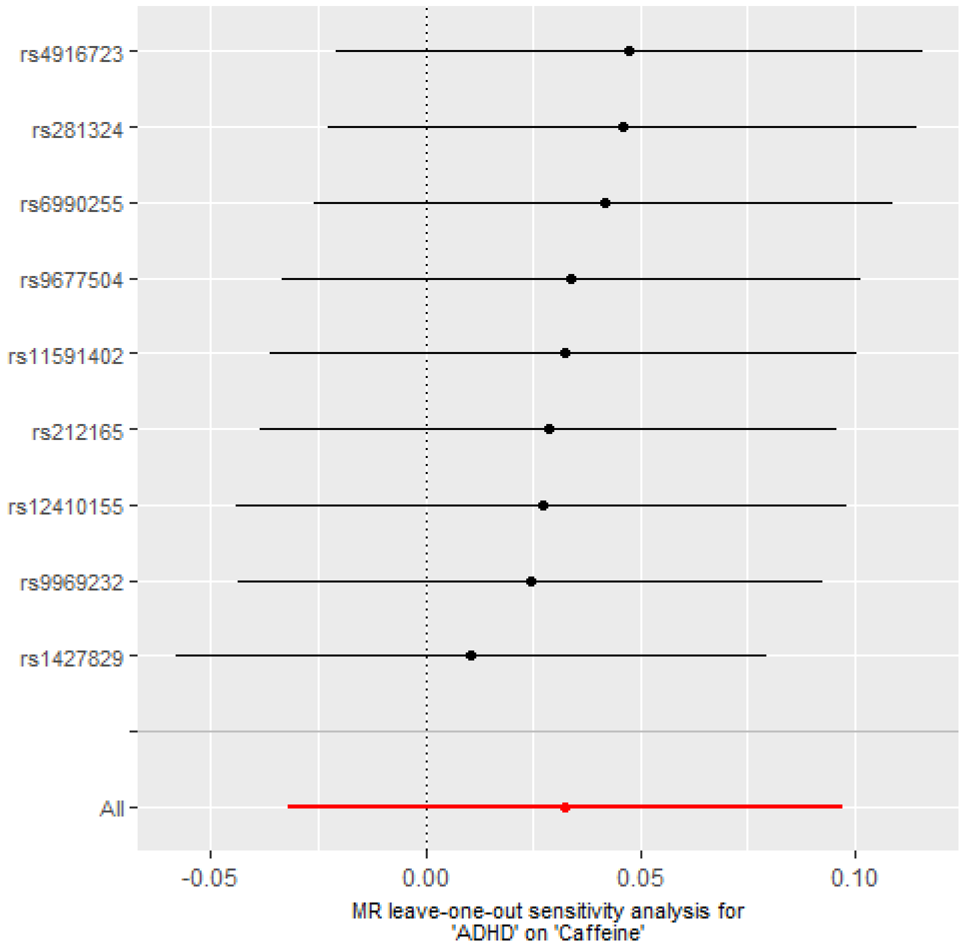

**Supplementary Figure 10.** Leave-one-out analyses depicting the results of Inverse Variance Weighted (IVW) Mendelian randomization analyses from liability to smoking initiation to ADHD risk after excluding each of the genetic variants from the analysis, one at the time. This gives an indication of whether or not the overall effect is driven by one particular genetic variant.

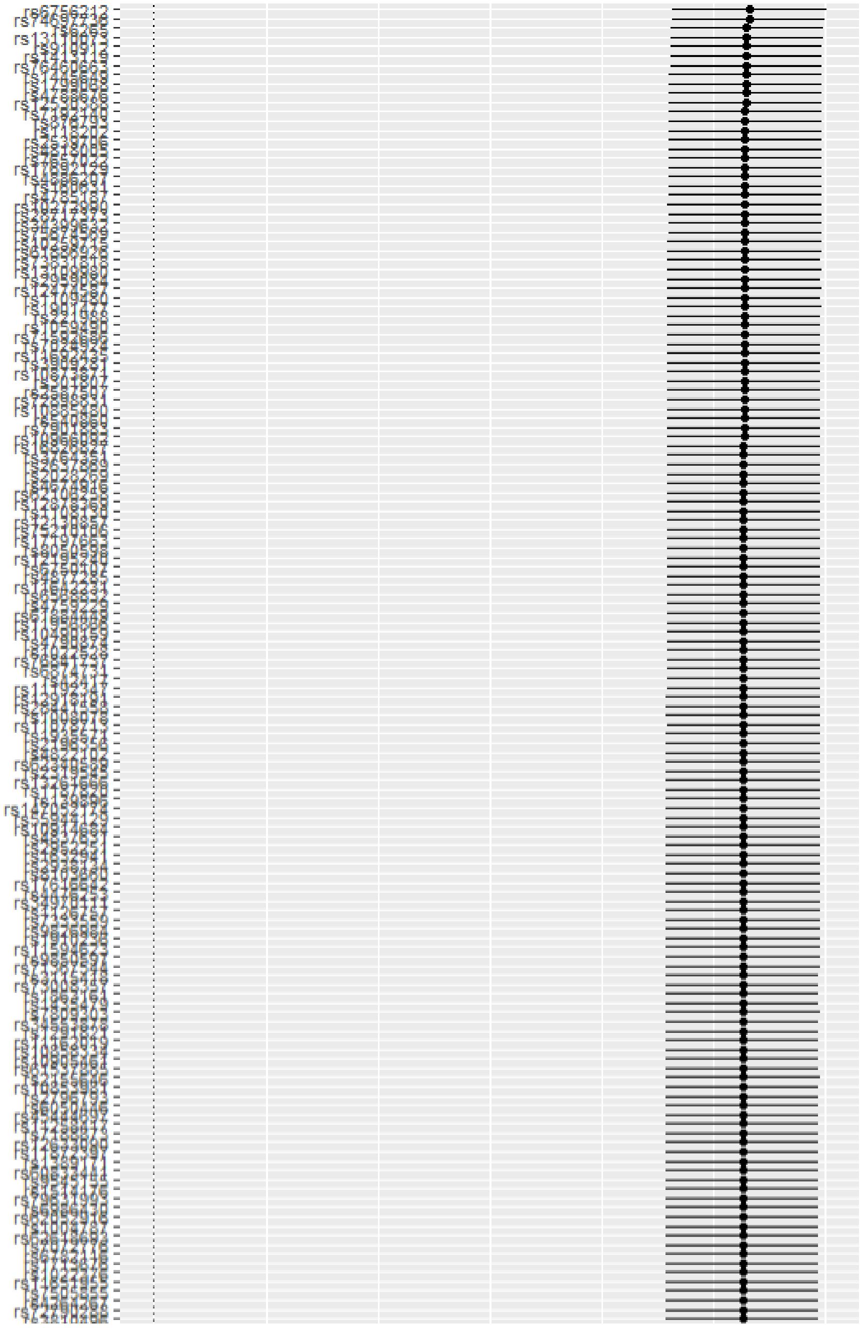

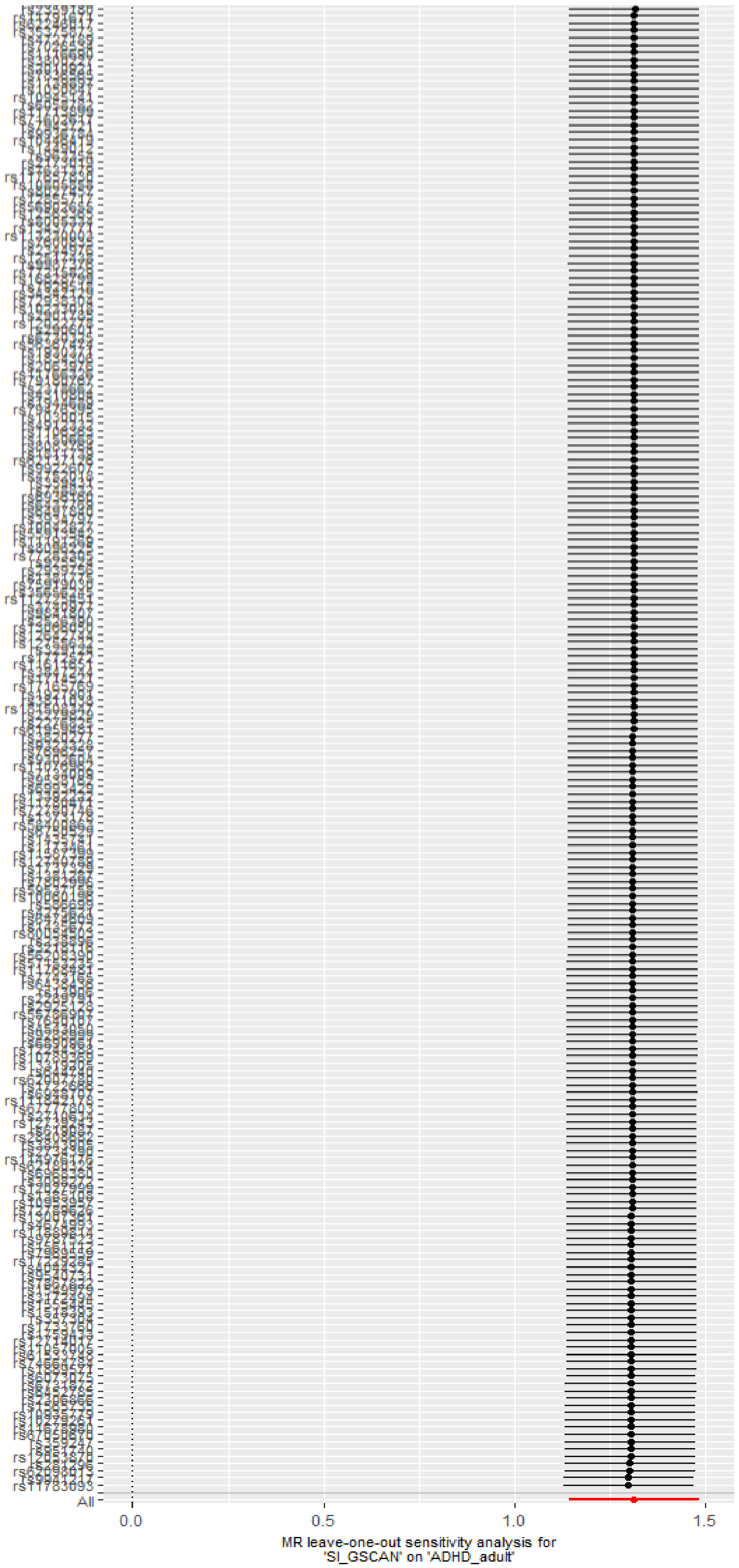
All
0.0
0.5
1.0
1.5
MR leave-one-out sensitivity analysis for
'SI_GSCAN' on 'ADHD_adult'

**Supplementary Figure 11.** Leave-one-out analyses depicting the results of Inverse Variance Weighted (IVW) Mendelian randomization analyses from alcohol drinks per week to ADHD risk after excluding each of the genetic variants from the analysis, one at the time. This gives an indication of whether or not the overall effect is driven by one particular genetic variant.

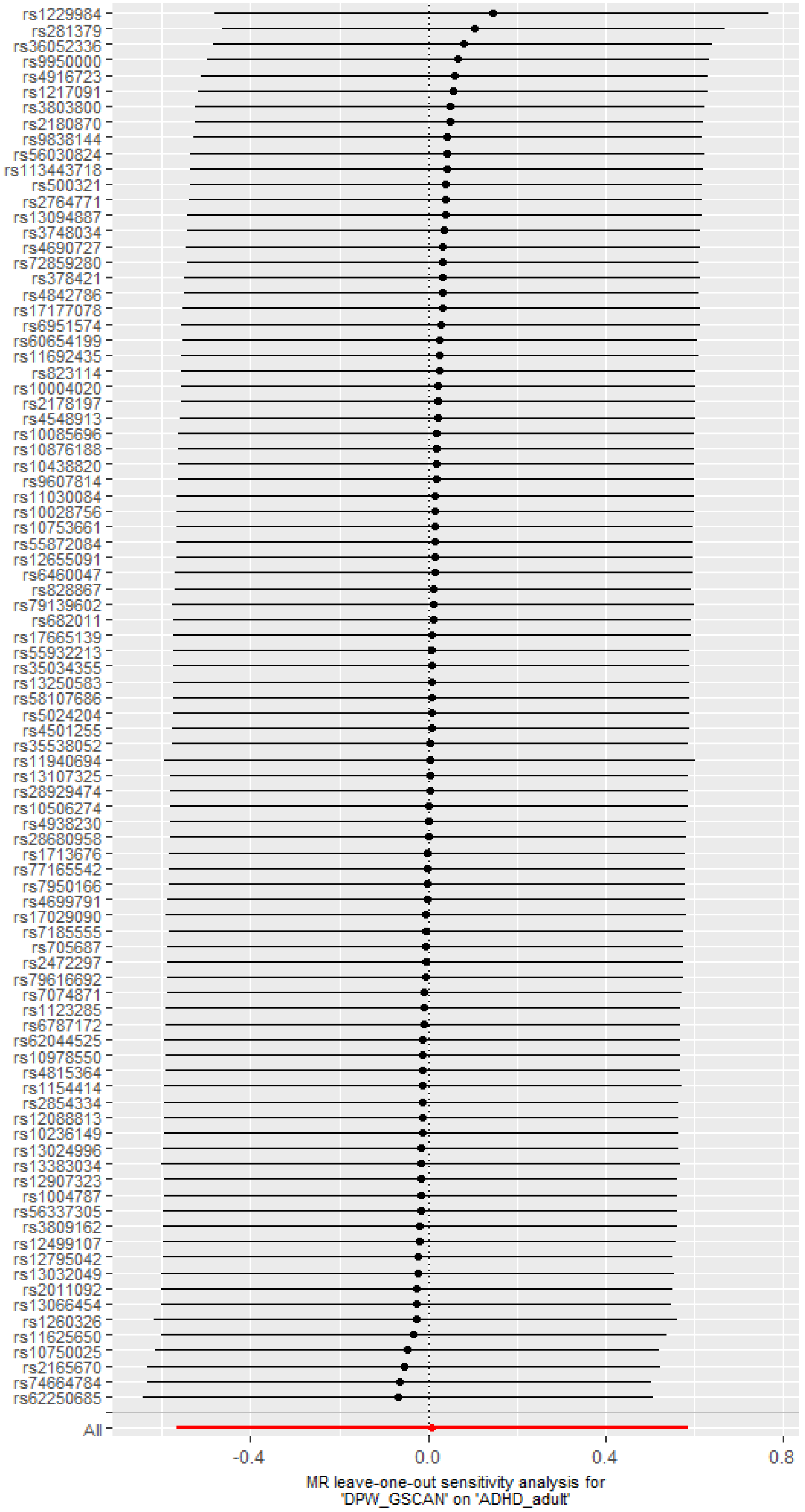

**Supplementary Figure 12.** Leave-one-out analyses depicting the results of Inverse Variance Weighted (IVW) Mendelian randomization analyses from liability to alcohol problems to ADHD risk after excluding each of the genetic variants from the analysis, one at the time. This gives an indication of whether or not the overall effect is driven by one particular genetic variant.

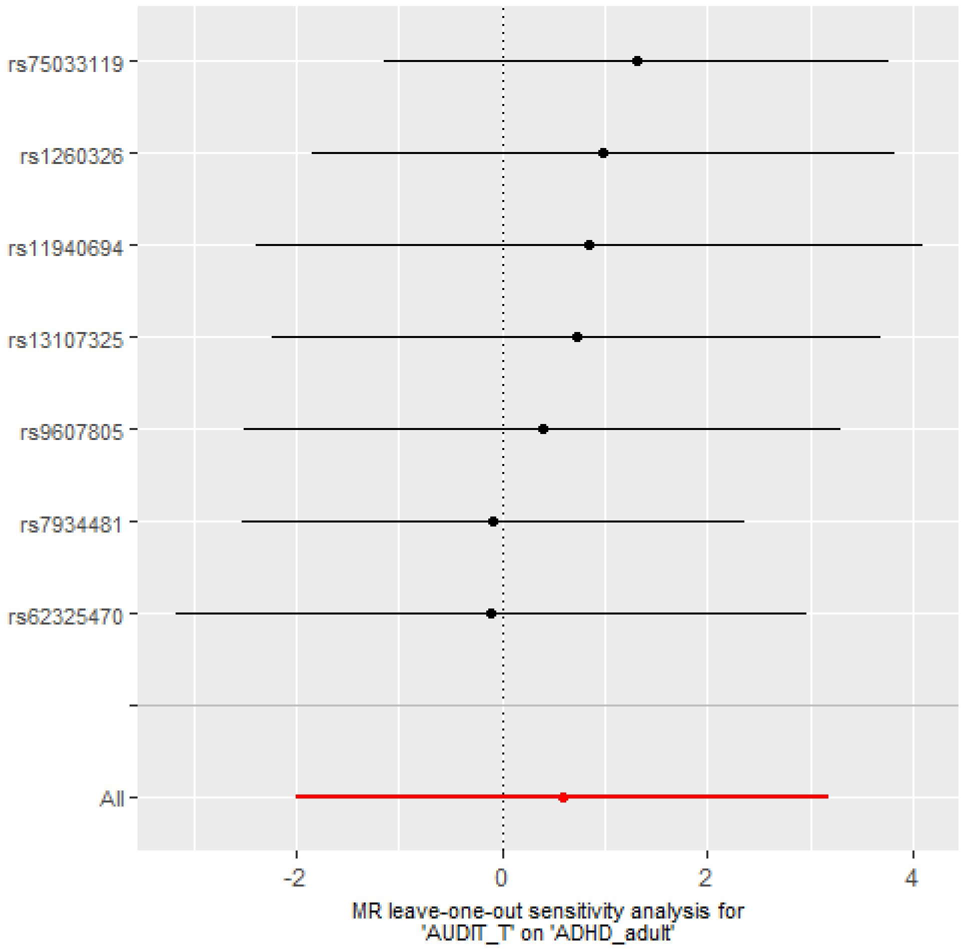

**Supplementary Figure 13.** Leave-one-out analyses depicting the results of Inverse Variance Weighted (IVW) Mendelian randomization analyses from liability to alcohol dependence to ADHD risk after excluding each of the genetic variants from the analysis, one at the time. This gives an indication of whether or not the overall effect is driven by one particular genetic variant.

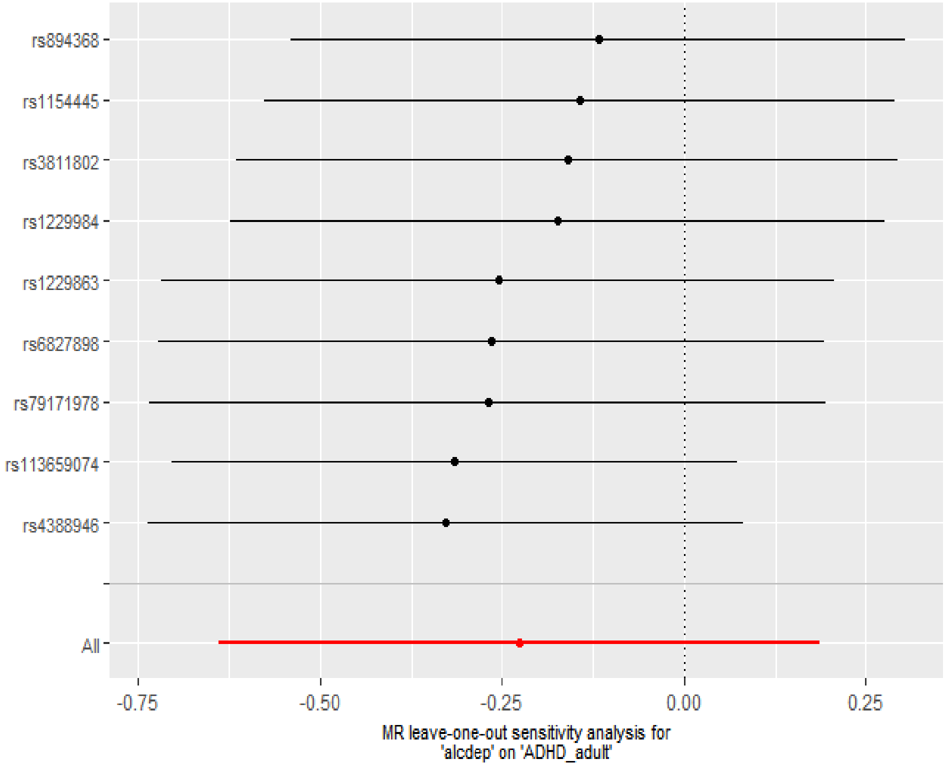

**Supplementary Figure 14.** Leave-one-out analyses depicting the results of Inverse Variance Weighted (IVW) Mendelian randomization analyses from liability to cannabis initiation to ADHD risk after excluding each of the genetic variants from the analysis, one at the time. This gives an indication of whether or not the overall effect is driven by one particular genetic variant.

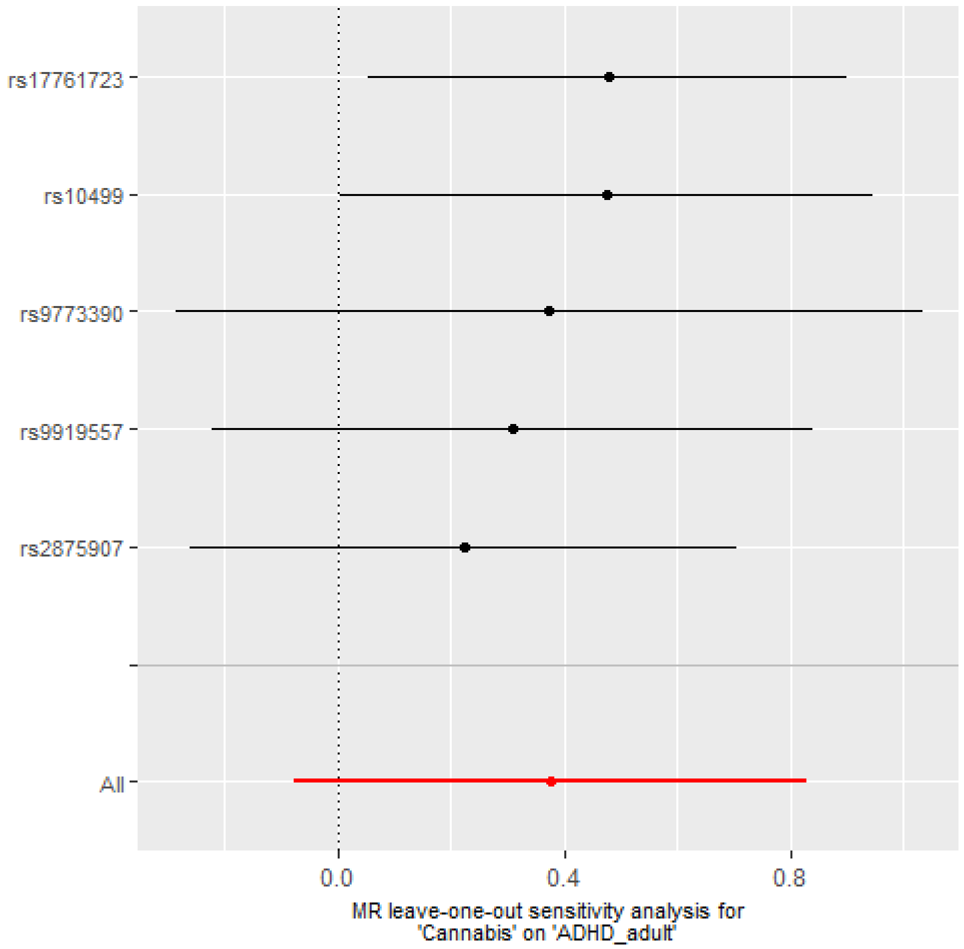

**Supplementary Figure 15.** Leave-one-out analyses depicting the results of Inverse Variance Weighted (IVW) Mendelian randomization analyses from cups of coffee per day to ADHD risk after excluding each of the genetic variants from the analysis, one at the time. This gives an indication of whether or not the overall effect is driven by one particular genetic variant.

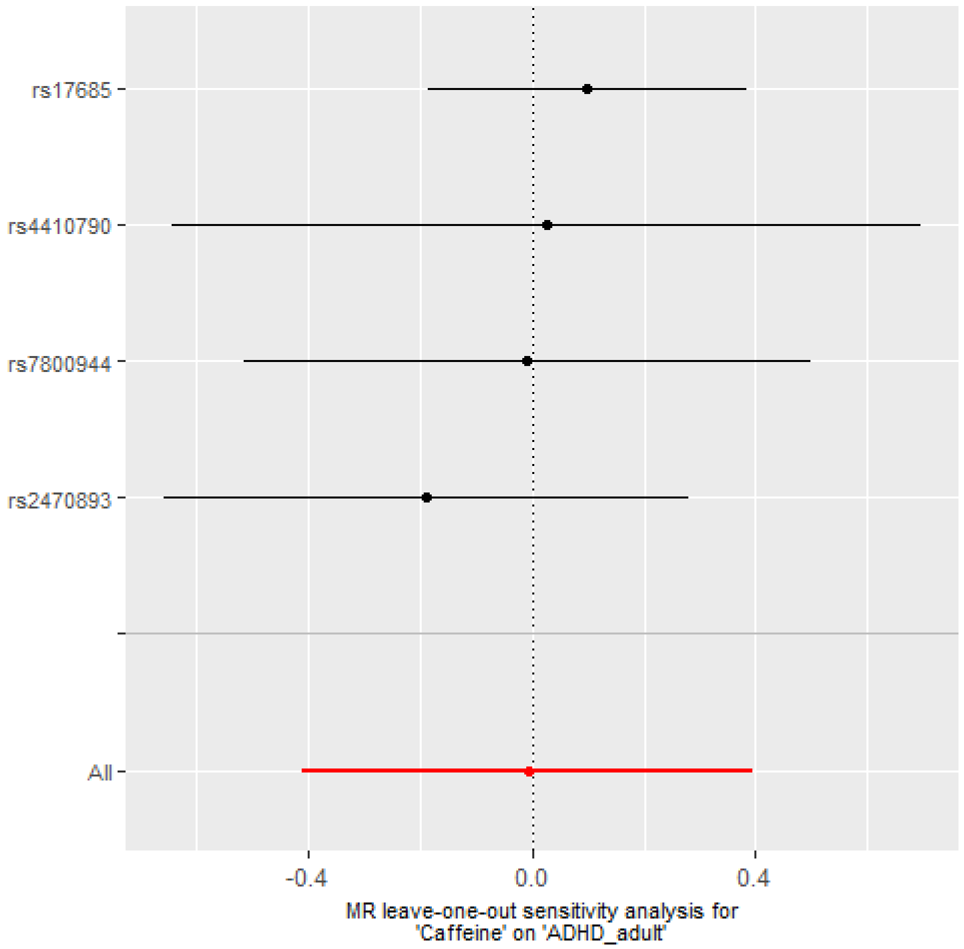
